# Patterning and folding of intestinal villi by active mesenchymal dewetting

**DOI:** 10.1101/2023.06.25.546328

**Authors:** Tyler R. Huycke, Hikaru Miyazaki, Teemu J. Häkkinen, Vasudha Srivastava, Emilie Barruet, Christopher S. McGinnis, Ali Kalantari, Jake Cornwall-Scoones, Dedeepya Vaka, Qin Zhu, Hyunil Jo, William F. DeGrado, Matt Thomson, Krishna Garikipati, Dario Boffelli, Ophir D. Klein, Zev J. Gartner

**Author notes:** Equal contribution (Tyler R. Huycke, Hikaru Miyazaki, Teemu J. Häkkinen).

## Abstract

Tissue folding generates structural motifs critical to organ function. In the intestine, bending of a flat epithelium into a periodic pattern of folds gives rise to villi, the numerous finger-like protrusions that are essential for nutrient absorption. However, the molecular and mechanical mechanisms driving the initiation and morphogenesis of villi remain a matter of debate. Here, we identify an active mechanical mechanism that simultaneously patterns and folds intestinal villi. We find that PDGFRA+ subepithelial mesenchymal cells generate myosin II-dependent forces sufficient to produce patterned curvature in neighboring tissue interfaces. At the cell-level, this occurs through a process dependent upon matrix metalloproteinase-mediated tissue fluidization and altered cell-ECM adhesion. By combining computational models with *in vivo* experiments, we reveal these cellular features manifest at the tissue-level as differences in interfacial tensions that promote mesenchymal aggregation and interface bending through a process analogous to the active de-wetting of a thin liquid film.

## INTRODUCTION

Many organs comprise functional epithelial sheets overlying a supportive mesenchymal stroma, connected through a common basement membrane interface. A critical step in organ morphogenesis and symmetry breaking is the folding of these tissue interfaces. Distinct tissues generate stereotyped folds, such as the cortical folds of the brain^1^, fingerprints and hair follicles of the skin^2^, branches of the mammary^3^ and clefts of the salivary gland^4^, bronchioles of the lung^5, 6^, and crypts and villi of the intestine^7–10^. Each of these folding patterns serves a function in the adult organ. Interfacial folding can arise through active forces, such as apical/basal epithelial constriction, or passive forces, such as mechanical instabilities produced from differential growth in tissue composites^11^. How interfacial folding occurs robustly across diverse contexts and is mechanistically coupled with tissue patterning to generate complex, functional, and morphologically diverse tissues is a fundamental question with implications for tissue engineering, developmental biology, and regenerative medicine.

In the embryonic small intestine, the emergence of positive curvature at the epithelial-mesenchymal interface demarcates the sites of forming villi, the elongated protrusions that expand the absorptive surface area of the gut nearly 100-fold relative to an otherwise non-folded surface geometry^12^. Postnatally, an additional round of tissue folding, this time generating negative curvature, leads to the formation of crypts that house intestinal stem cells essential for tissue homeostasis and regeneration^7^. In addition to establishing the repetitive and stereotyped architecture that enables digestive functions, the geometry of these folds specifies epithelial cell identities. For example, tissue folding leads to the reshaping of morphogen gradients that restricts stem and progenitor cells to the intervillus space, while zones of non-proliferation and differentiated cell types become localized to the emerging villi^13^. This stereotyped tissue geometry also determines cell fate directly by specifying stem cells to localize at regions of high curvature^14–16^. The zonation of cellular function, first established during development, is maintained into adulthood and is critical for homeostasis and regeneration^17^.

Tissue folding emerges at the earliest stages of villus formation and is proposed to occur through species-specific modes of morphogenesis^18^. In avians, bending of the luminal surface progresses through a series of elastic buckling steps mediated by the sequential differentiation of orthogonally aligned smooth muscle layers that restrict the expansion of the faster-growing inner mesenchyme and epithelium^8^. Compressive stresses are relieved by transforming the luminal surface into a series of longitudinal ridges that are then compressed further to zigzags, from which villi subsequently arise. In contrast, mammalian villi emerge directly from a flat luminal surface wherein tissue folding coincides spatiotemporally with the appearance of subepithelial mesenchymal aggregates, termed villus clusters (**Figure 1A**). Villus clusters have been hypothesized to be either a cause or consequence of tissue folding; arising by a Turing-like chemical reaction diffusion system to pattern where villi form, or arising due to the specifying actions of concentrated morphogens from an already folded epithelium^9, 10, 19, 20^. Whether villus clusters are actively involved in the generation of curvature remains to be determined, and more generally, the physical and molecular mechanism that establishes patterns of curvature in the mammalian intestine remains unclear.

**Figure 1.**
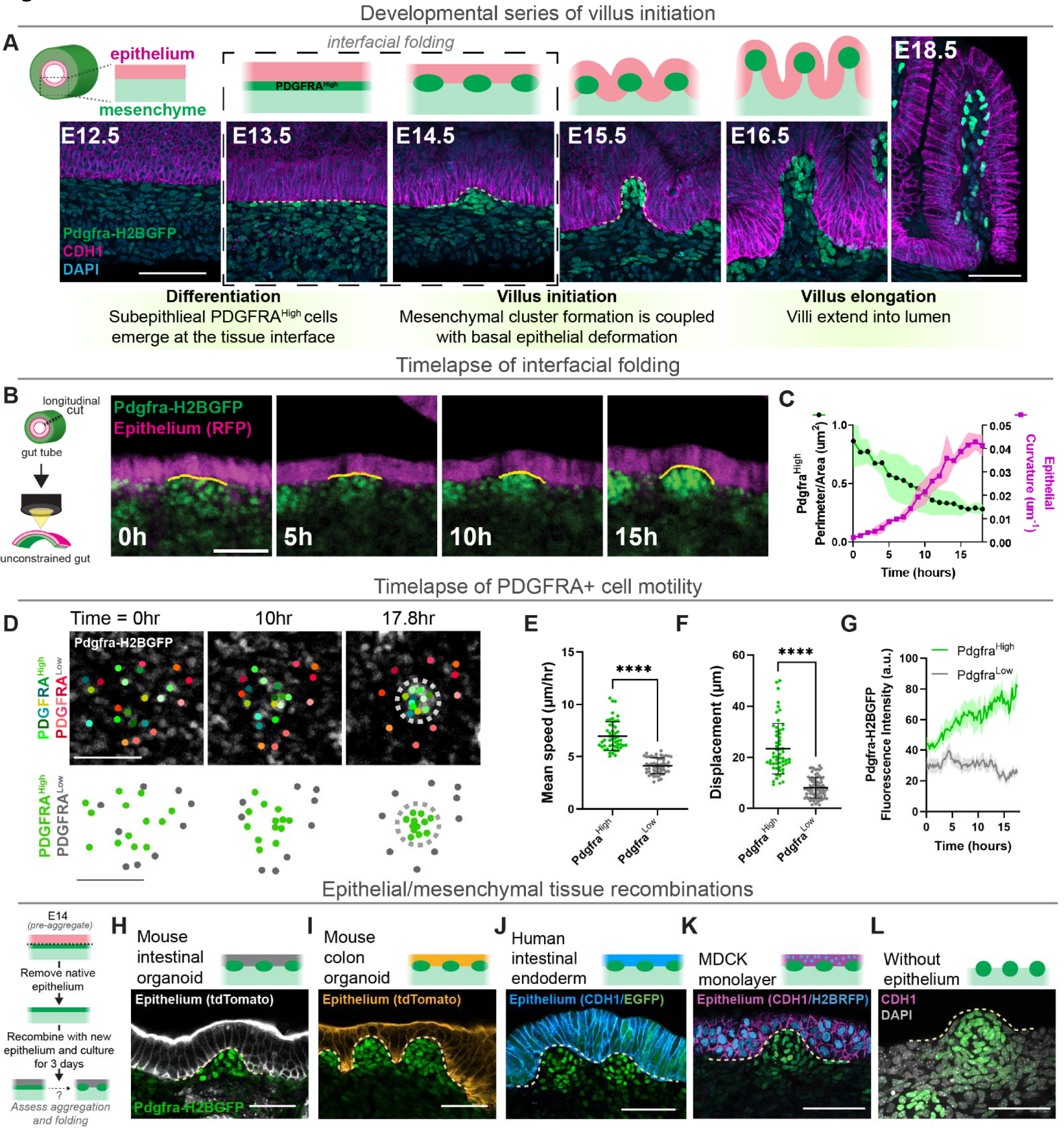
PDGFRA^High^ cell aggregation is sufficient to drive interfacial folding and initiate villi. (A) Optical sections showcasing key developmental stages of intestinal villus morphogenesis from *PDGFRA^H2BGFP^* mice, immunostained for CDH1 (E-Cadherin) to label the epithelium. Highlighted box emphasizes the emergence of patterned interfacial folding, indicated by a dotted yellow line at the epithelial-mesenchymal boundary. (B) Snapshots from a timelapse video of an E14 intestinal explant from *Pdgfra^H2BGFP^; Shh^Cre^; R26R^tdTomato^* mice showing the co-emergence of mesenchymal aggregates and interfacial curvature, highlighted with a yellow line. (C) Quantification of PDGFRA^High^ aggregate perimeter/area ratio and interfacial curvature from timelapse videos as in (B). Curvature measurements were taken at 1 hour intervals. Error bars are mean ± SD. *n*=5 aggregates measured from two explants. (D) Example of cells tracked in a local region during the aggregation process from timelapse videos. Cells are color coded by identities (red shades = PDGFRA^Low^, non aggregating cells; green shades = PDGFRA^High^, aggregating cells. Bottom panels represent cell tracks in top panels. (E) Quantification of mean speed of tracked nuclei from timelapse movies. Each point represents the average speed calculated for a single nucleus across a 20 hour timeframe, collected from *n = 3* intestine explants. (F) Quantification of displacement of tracked nuclei from timelapse movies. (G) Fluorescence intensity of *PDGFRA^H2BGFP^* reporter over the course of a timelapse in each lineage. Collected from *n =* 15 cells for each phenotype (PDGFRA^Low^ or PDGFRA^High^) from three intestinal explants (H-L) Optical sections from recombinant tissues, color coded by origin of epithelia for (H) tdTomato-labeled adult mouse enteroid epithelium, (I) tdTomato-labeled adult mouse colonoid epithelium, (J) cytoplasmic GFP-labeled HIO-derived endoderm, (K) nuclear RFP-labeled MDCK cells, and (L) no epithelium (negative CDH1 stain). Dashed lines demarcate the folded epithelial-mesenchymal interfaces at sites of mesenchymal aggregation. Recombinant tissues in panels K and L are incubated in the presence of 1µM SAG. Schematic to the left of panel H depicts the experimental setup. All images are representative of at least 9 tissue recombinants per condition. Scale bars = 50 µm; \*\*\*\**p<0.0001*

Here, we provide evidence that the patterning and folding of the small intestine epithelial-mesenchymal interface occur by a distinct and surprising mechanism involving the dynamic mechanical properties of villus clusters. We first show that the cellular activities leading to mesenchymal cell aggregation are sufficient to drive interfacial folding. Then, by integrating theory and experiment, we demonstrate that the mesenchyme behaves as an active fluid whose morphology is dictated by surface tensions that emerge at its interface with surrounding tissue layers.

## RESULTS

### Epithelial folding of the mouse intestine is coupled to mesenchymal aggregation

Previous work suggests that, in contrast to the chick, global compressive forces provided by smooth muscle layers are dispensable for initiating tissue curvature in the mouse^9^. We confirmed and expanded upon these findings to demonstrate that in the absence of functional muscle, global compressive forces or differential growth, the PDGFRA-expressing mesenchymal aggregates that comprise villus clusters still arise directly at the sites where curvature is generated in the epithelium (**Figure S1A-C**) (**Figure 1A**). These observations support an alternative model wherein aggregating mesenchymal cells situated at the epithelial-mesenchymal interface are the cause of the interfacial folding that initiates villus formation^9, 10, 12, 18, 19^, rather than the consequence of folding as occurs in the chick^13, 21^. To gain deeper insight into this problem and to determine which arises first – mesenchymal aggregates or tissue curvature – we devised a system to perform long-term timelapse imaging of isolated whole gut tissue explants, which differentiate and undergo morphogenesis *ex vivo* in a manner nearly identical to *in vivo.* This system enables imaging across long time periods (>24 hours) and at high temporal resolution, allowing us to capture morphogenesis of both the mesenchymal (marked by *PDGFRA^H2BGFP^*) and epithelial (marked by *Shh^Cre^; R26R^tdTomato^*) compartments as well as interfacial folding in considerably more detail than previous reports. We focused our study on the critical symmetry breaking event that occurs between embryonic days (E)13.5 and E14.5 when patterning and folding emerge from an initially undifferentiated and unfolded tissue interface (highlighted in **Figure 1A**). We observed that the emergence of epithelial curvature and the aggregation of mesenchymal cells expressing high levels of PDGFRA (henceforth PDGFRA^High^ cells) into villus clusters occurs simultaneously (**Figure 1B, 1C, and Video S1**). The tight coupling of these events is consistent with tissue curvature arising from the activities of cells in either the epithelium, the mesenchyme, or coordinated by both.

To better distinguish among these possibilities, we used intestinal explant imaging to examine the underlying epithelial and mesenchymal cell dynamics. While epithelial cells show minimal movement and no neighbor exchange (**Figure S2A**), the upper (subepithelial) layer of mesenchyme is highly motile, consistent with previously reported short-term timelapse data^22^ and inconsistent with a reaction-diffusion dominated mechanism where one would expect to observe patterned, local differentiation of PDGFRA^High^ foci^9^ (**Video S2**). Direct quantification of individual cell movements showed that PDGFRA^High^ cells move into clusters from distal locations, exhibiting higher motility than neighboring PDGFRA^Low^ cells that do not form aggregates, and gradually express higher levels of PDGFRA over time (**Figure 1D-G**). Thus, motility-driven PDGFRA^High^ mesenchymal aggregation and the generation of tissue curvature within the overlying static epithelial interface are tightly linked, prompting us to further probe whether folding might be driven directly by the mesenchyme.

### Mesenchymal aggregation drives interfacial folding to initiate villus formation

To conclusively determine the specific source of cellular activities that generate interfacial curvature, we asked whether the mesenchyme is sufficient to initiate tissue folding independent of the overlying surface identity. We therefore performed a series of tissue recombination experiments wherein we non-enzymatically removed the native intestinal epithelium at E14, just prior to the onset of mesenchymal aggregation, and replaced it with epithelia from other sources. First, we recombined the mesenchyme with epithelia of organoids derived from the adult mouse intestine or colon (which does not normally contain villi or mesenchymal aggregates) that had been propagated *in vitro* in the absence of mesenchyme. In both cases, PDGFRA^High^ mesenchymal cell aggregation and epithelial deformation occured (**Figure 1H and 1I**), although the final pattern and geometry of folding in each case was distinct (discussed below). Next, we differentiated human intestinal organoids (HIOs) from pluripotent stem cells^23^, isolated their endodermal layers as epithelial sheets, and recombined these with the mouse mesenchyme. Again, mouse PDGFRA^High^ mesenchymal cells formed aggregates, and folds emerged in the overlying epithelium, now human-derived, suggesting that an ability to fold the surface is not species-specific and that the human gut might form villi in the same manner as in the mouse (**Figure 1J**). Indeed, we observed similar co-occurrence of mesenchymal aggregates and tissue folding in transplanted human intestinal organoids (tHIOs) (**Figure S2B**). We then examined whether mesenchymal cell aggregation could generate patterned folds in a monolayer of Madin-Darby canine kidney (MDCK) cells, an immortalized cell line derived from a completely different organism, tissue of origin, and embryonic germ layer than the murine intestinal endoderm. However, in basal media conditions the mesenchyme degenerated, failed to aggregate, and no curvature formed in the MDCK layer (**Figure S2C and S2D**). We reasoned that this outcome could be due to a lack of Hedgehog (Hh) ligands, which are expressed by the intestinal epithelium, but not highly by MDCK cells^24, 25^, and which signal in a paracrine fashion to the mesenchyme to regulate villus formation^19, 26, 27^. Indeed, treatment of recombinant mesenchyme-MDCK tissues with the Hh pathway activator Smoothened Agonist (SAG) enabled tissue viability and was sufficient to induce mesenchymal aggregation and folding of the MDCK monolayer (**Figure 1K and S2D**). The ability of the mesenchyme to fold the MDCK monolayer prompted us to test whether the mesenchyme alone could generate interfacial folds. Strikingly, PDGFRA^High^ cells aggregated at the surface of epithelia-free SAG-treated mesenchyme slabs, and these aggregates generated characteristic curvature at the tissue-media interface (**Figure 1L**). Thus, the subepithelial mesenchyme is sufficient to drive interfacial folding in the absence of an epithelium, and tissue surface curvature is therefore generated directly by the aggregation of motile PDGFRA^High^ mesenchymal cells.

### Emergence of distinct subepithelial mesenchymal populations during development

Having determined that subepithelial PDGFRA^High^ mesenchymal cell aggregation generates the curvature necessary to initiate intestinal villi, we set out to identify the molecular and mechanical mechanisms through which these cells aggregate and generate patterned interfacial folds. We turned to single-cell RNA sequencing to identify transcriptional differences between PDGFRA^High^ cells and their non-aggregating PDGFRA^Low^ neighbors that could provide clues to how these cells initiate morphogenesis at the tissue interface. We gathered an integrated data set from intestinal tissues isolated from five developmental stages spanning the timeframe of villus initiation (E13.5, E14.5, E15.5, E16.5, and E18.5) using the MULTI-seq approach^28^ and enriched for PDGFRA^High^ cells by Fluorescence-Activated Cell Sorting (FACS) (**Figure S3A-C**).

We identified six major cell types within the embryonic mouse intestine, including epithelial, mesenchymal, endothelial, immune, mesothelial and enteric nervous system (ENS) populations (**Figure S3D and S3E**). We then subsetted mesenchymal cell types based on well-established markers and subsequently the subepithelial mesenchyme based on its expression of *Bmp4*, a marker of mesenchymal cells situated between the epithelium and circumferential muscle layer^9, 26, 27^ (**Figure S3F and S3G**). PDGFRA^High^ expressing cells were highly enriched within this subepithelial mesenchymal cell group, as expected (**Figure 2A and S3G**). When the FACS-sorted PDGFRA^High^ and PDGFRA^Low^ populations from this subsetted population are visualized by Uniform Manifold Approximation and Projection (UMAP), we observed differentiation trajectories for each population spanning the key developmental stages of villus morphogenesis (**Figure 2B and S3J**). Furthermore, cell clustering analysis of the subepithelial mesenchyme revealed the presence of presumptive cycling progenitor populations as well as the PDGFRA^High^ and PDGFRA^Low^ lineages, with clusters tracking closely with the progression of developmental timepoints (**Figures 2B, 2C and S3I-K**). These subepithelial clusters express unique marker genes (**Supplementary Table 1**) including those previously associated with *PDGFRA^High^*subepithelial mesenchymal populations in the human intestine (e.g. *F3* and *Dll1*), as well as genes expressed by adult PDGFRA^High^ subepithelial myofibroblasts/telocytes (e.g. *Foxl1*) and PDGFRA^Low^trophocytes (e.g. *Rspo2/3*), known for their roles in establishing the intestinal stem cell niche (**Figure S3H**)^17, 29–31^.

**Figure 2.**
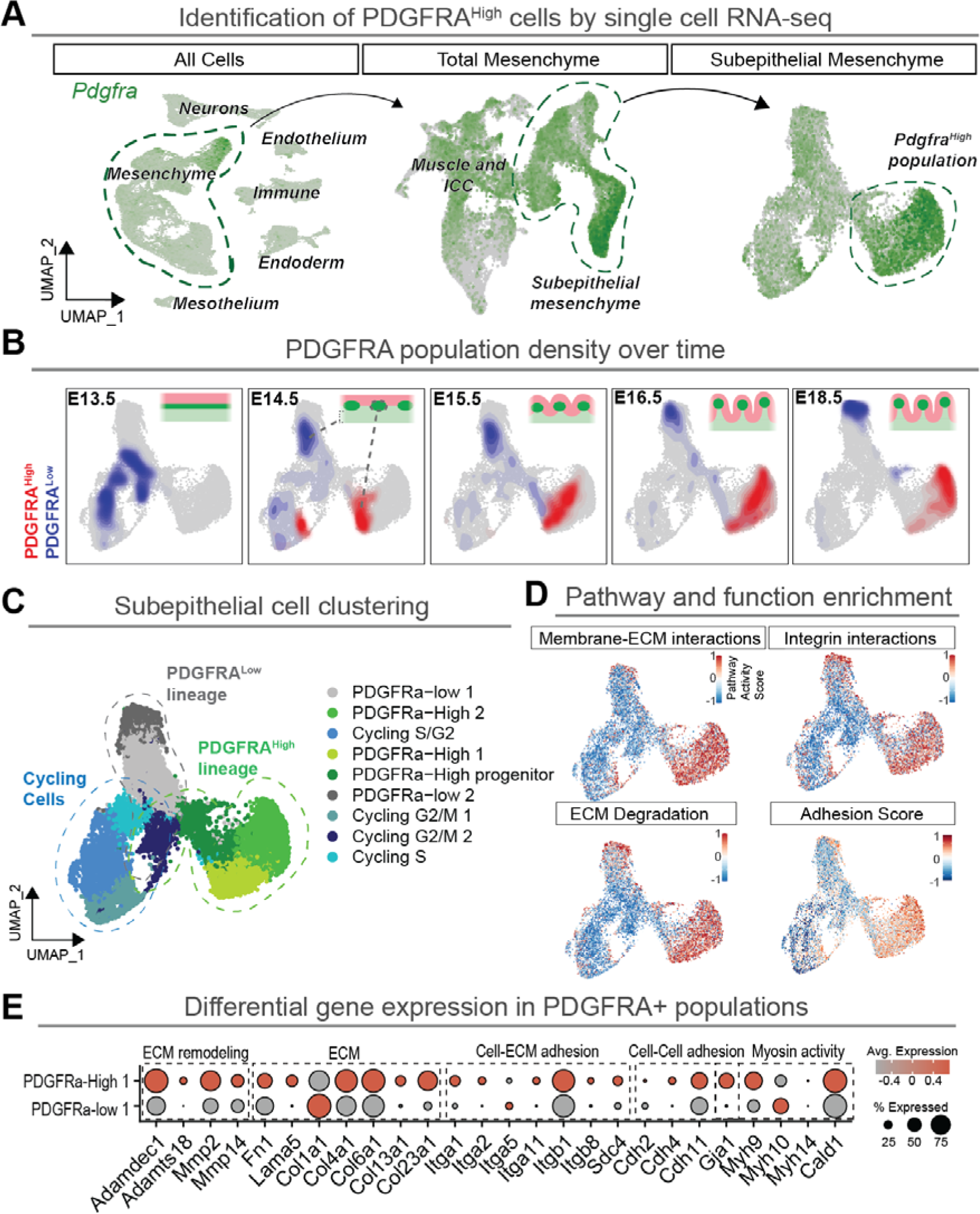
PDGFRA^High^ cells upregulate expression of ECM remodeling, adhesion, and contractile programs. (A) Uniform Manifold Approximation and Projection (UMAP) dimensionality reduction of scRNA-seq data from all cells collected at E13.5, E14.5, E15.5, E16.5, and E18.5, then subset and re-clustered for the mesenchyme alone and then the subepithelial mesenchyme, which excludes smooth muscle and interstitial cells of Cajal lineages. (B) Density plots showing the location of the PDGFRA^High^ population (cells enriched for by FACS) and the PDGFRA^Low^ population over each developmental stage collected to highlight the divergence of the two lineages and stage-specific locations. (C) Cell clusters in the PDGFRA+ subepithelial subset. (D) UMAPs of gene modules contributing to cell-cell and cell-matrix adhesion identified by the Reactome Database and plotted as a Reactome Pathway Activity score calculated by AUCell and the Adhesion Score defined by cell adhesion interacting genes (See Methods). (E) DotPlot comparing expression for select key genes related to noted cell and molecular functions between the PDGFRA-High 1 and PDGFRA-low 1 clusters.

### Aggregate-forming PDGFRA^High^ cells express a unique cell-cell and cell-ECM interaction transcriptional program

To understand the cellular and molecular programs that underlie formation of PDGFRA^High^ mesenchymal aggregates, we first performed differential gene expression (DGE) analysis comparing PDGFRA^High^ cells (PDGFRA-High 1 cluster) and PDGFRA^Low^ subepithelial cells (PDGFRa-low 1 cluster) (**Supplementary Table 2**). Pathway enrichment for PDGFRA^High^ marker genes using Gene Ontology (GO), Kyoto Encyclopedia of Genes and Genomes (KEGG), and the Reactome database revealed that PDGFRA^High^ cells are significantly enriched for programs related to cell adhesion and cell-ECM interactions (**Figure 2D, S3M-P and Supplementary Table 3**). Furthermore, consensus non-negative matrix factorization (cNMF)^32^ and Waddington-OT (WOT) analysis^33^ identified candidate driver genes, broadly overlapping with the above cluster-based DGE analysis, that become specifically activated in a coordinated manner during PDGFRA^High^ differentiation and are uniquely upregulated in PDGFRA^High^ vs. PDGFRA^Low^ lineages (**Figure 2E, S3Q, S3R and Supplementary Tables 4 and 5**). Together, these transcriptional analyses of the subepithelial mesenchyme point to a mechanistic role for actomyosin activity (e.g., *Myh9, Myh10*), cell-cell (e.g., *Cdh4, Cdh11*) and cell-ECM adhesion (e.g., *Itga2, Col6a1, Col23a1*), and matrix remodeling (e.g., *Mmp2, Mmp14)* in PDGFRA^High^ mesenchymal cell aggregation (**Figure 2E**).

### Non-muscle myosin activity in PDGFRA^High^ cells drives cell motility, aggregation, and villus formation

To functionally interrogate the role of these upregulated cellular programs in mesenchymal aggregation and interfacial folding, we began by examining the spatial distribution of key markers for non-muscle myosin activity at the protein level. Consistent with transcriptional analysis, MYH9 (NMIIA) is enriched specifically in aggregating PDGFRA^High^ cells, MYH10 (NMIIB) is expressed more broadly throughout the mesenchyme, and MYH14 (NMIIC) expression is mostly restricted to the epithelium (**Figure 3A and 3B**). Moreover, expression of phospho-myosin light chain (pMLC), a marker for actomyosin activity, and the upstream Myosin Light Chain Kinase (MLCK) are upregulated in villus clusters relative to the epithelium and PDGFRA^Low^ mesenchyme, further confirming that the PDGFRA^High^ mesenchyme actively generates forces to produce interfacial curvature independently of the epithelium (**Figure 3A, 3B, and S4A**).

**Figure 3.**
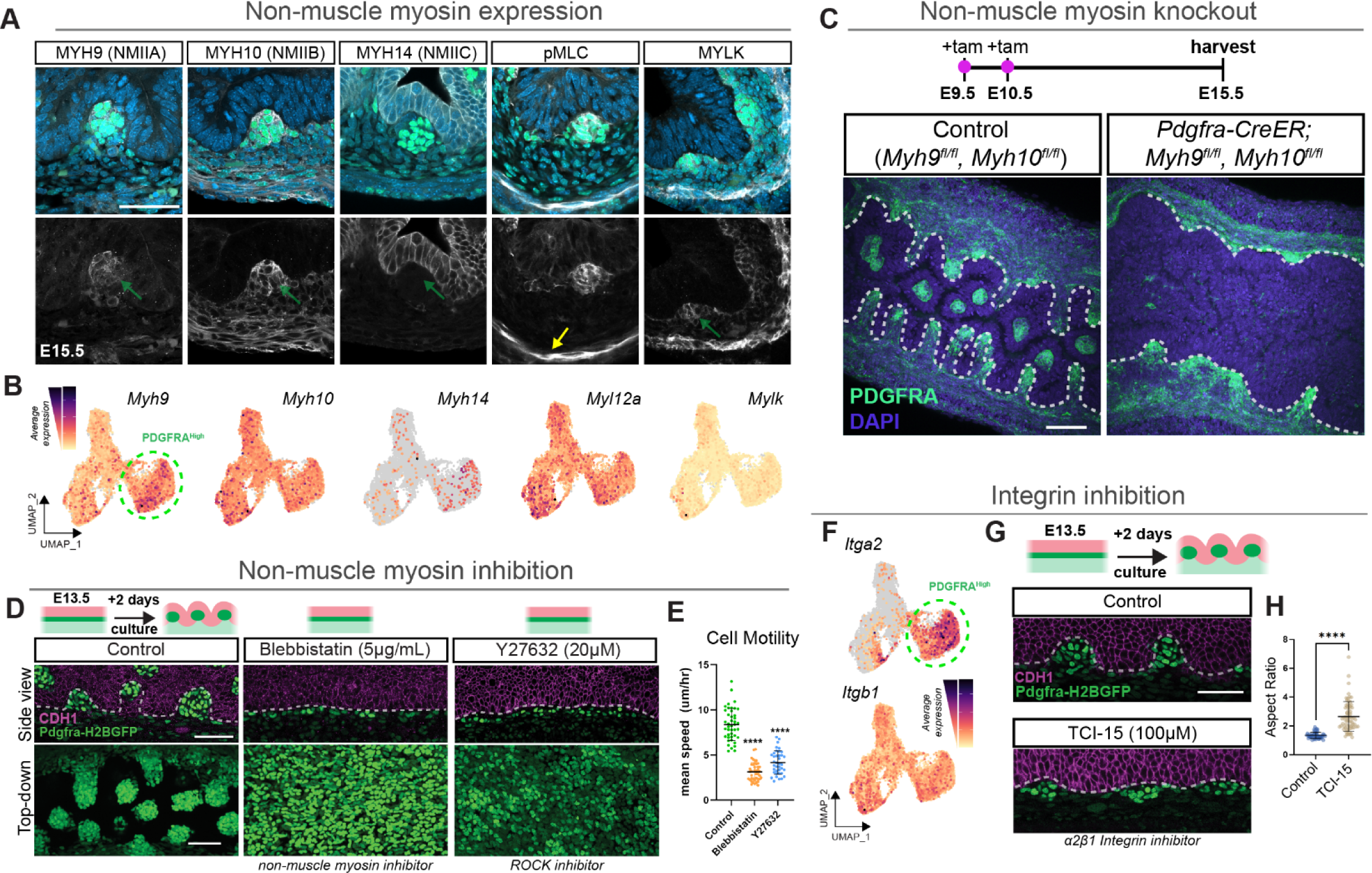
Mesenchymal non-muscle myosin activity and integrin-mediated adhesion are required for cell aggregate compaction and villus initiation. (A) Immunofluorescence (IF) in cryosections of E15.5 mouse small intestines stained for various actomyosin activity markers. Green arrows mark the location of the mesenchymal aggregates in bottom single channel images, and yellow arrow denotes expected expression of pMLC in the smooth muscle layer. (B) UMAP plots showing RNA expression for protein products labeled by IF in (A). Region circled denotes the PDGFRA^High^ population as determined in Figure 2. (C) Optical sections from whole mount IF of the intestines at E15.5 from a control or conditional mutant in which non-muscle myosin (*Myh9* and *Myh10*) were knocked out in *Pdgfra*-expressing cells of the mesenchyme. Optical sections are from the midline of the intestine in the proximal jejunum of the small intestine. Dotted white lines denote the epithelial-mesenchymal interface. Images are representative of *n =* 9 double mutants. (D) Optical sections from whole mount IF showing side view (tissue cross section) or top-down views (*en face*) of the subepithelial mesenchyme. Tissues are explants grown ex vivo from E13.5 for 2 days and treated with noted inhibitors. Images are representative of *n* ≥ *6* samples per condition. (E) Quantification of cell motility (mean speed) from timelapse movies of explants treated with inhibitors as in panel D. Timelapse movies were acquired by culturing E14 guts and imaging middle (jejunum) regions of the intestine that had not begun aggregating at the start of the movie. Error bars are mean ± SD. Each point represents a cell, measured from *n = 3* tissues per condition. (F) UMAP plots of Integrin expression in the subepithelial mesenchyme. (G) Optical sections from whole mount IF showing subepithelial interface of explants treated with the A2B1 integrin inhibitor TCI-15. (H) Quantification of aggregate aspect ratio from explants in panel G. Aggregates were measured across 4 samples per treatment. Error bars are mean ± SD, each point represents a distinct aggregate measured across *n =* 5 treated or untreated explants. Scale bars = 50 µm; \*\*\*\**p<0.0001*

Next, we tested whether non-muscle myosin activity is required for mesenchymal aggregation and villus initiation using *Pdgfra^CreER^*to conditionally delete *Myh9* and *Myh10* in the *Pdgfra*-expressing mesenchyme. Conditional *Myh9/Myh10* double knockouts failed to form the typical periodic pattern of mesenchymal aggregates, with interfacial folding largely absent at E15.5, a time when villi are well formed in controls (**Figure 3C**). Conversely, PDGFRA^High^ cells still aggregated and generated interfacial folds when *Myh9/10* was conditionally deleted in the epithelium, despite epithelial dysplasia (**Figure S4B**). To determine how mesenchymal non-muscle myosins control the cell dynamics that enable clustering, we treated intestinal explants with the non-muscle myosin inhibitor blebbistatin or the ROCK inhibitor Y27632. In both cases, PDGFRA^High^ cells differentiated at the tissue interface (as noted by the increased expression of PDGFRA) but failed to move, aggregate and deform the interface (**Figures 3D, 3E**). Importantly, the effect of blebbistatin is reversible upon washout (**Figure S4C**), and therefore, molecular differentiation of PDGFRA^High^ cells versus tissue morphogenesis at this interface can be uncoupled by modifying the active mechanical properties of these cells. Thus, actomyosin activity within PDGFRA^High^ cells, mediated by MYH9 and MYH10, drives differential cell motility and cell aggregation to form villus clusters.

### Integrin-mediated cell interactions are required for proper mesenchymal aggregation and compaction

Having found that myosin is the motor that generates the forces driving cell aggregation, we next asked what molecular machinery engages with the motor to bind PDGFRA^High^ cells together into a compact aggregate at the epithelial interface. Our transcriptional analyses suggested that cell-ECM^34^ adhesion could be important for cohesion and compaction of PDGFRA^High^ aggregates. We hypothesized that these processes could be mediated specifically by ITGA2, which forms heterodimers with ITGB1 to bind various collagens, is uniquely upregulated at the RNA level within the PDGFRA^High^ cell lineage (**Figured 2E and 3F**), and, along with prospective binding partners *Col23a1* and *Col6a4*, is a top driver gene of the PDGFRA^High^ state identified by WOT and cNMF analyses (**Supplementary Tables 4 and 5**). We therefore tested integrin alpha 2 beta 1 (ITGA2B1) function in villus initiation by blocking its interaction with collagen using the inhibitor TCI-15^35^ in explant culture. ITGA2B1 inhibition led to strikingly altered aggregate morphology, with reduced compaction, greater and more variable aspect ratios, and a near absence of bending at the overlying epithelial interface (**Figures 3F-H and S4D**). While we cannot rule out a role for CDH4 and CDH11, two cadherins upregulated in PDGFRA^High^ aggregates (**Figure 2E and S4E**), these data indicate that ITGA2B1-mediated cell-ECM interactions are necessary for aggregate compaction, regular patterning, and interfacial folding.

### Loss of cell and ECM anisotropy precedes the onset of mesenchymal aggregation

The impact of ITGA2B1 perturbation combined with the upregulation of a cell-ECM interaction program at the transcriptional level prompted us to further characterize the nature of the cell-ECM interface during PDGFRA^High^ cell aggregation and the emergence of tissue curvature. We began by examining cell and ECM morphology at the key developmental stages during which these events occur. As villus initiation proceeds in a proximal-to-distal wave along the length of the gut, two steps of the process can be captured within a single developmental stage at E14.5, when aggregation is taking place in the proximal intestine but has not yet initiated in the distal intestine **(Figure 4A**)^19^. Examination of tissue architecture at this stage and within these regions revealed that cells and nuclei are aligned parallel to the long axis of the gut in the pre-aggregate distal regions, whereas this alignment has dissipated in the aggregate-initiating proximal gut. This pattern is mirrored by changes in the ECM: fibronectin and collagen fibers are aligned parallel to the proximal-distal axis in the distal gut but are not aligned in the proximal gut (**Figure 4A and S5A**).

**Figure 4.**
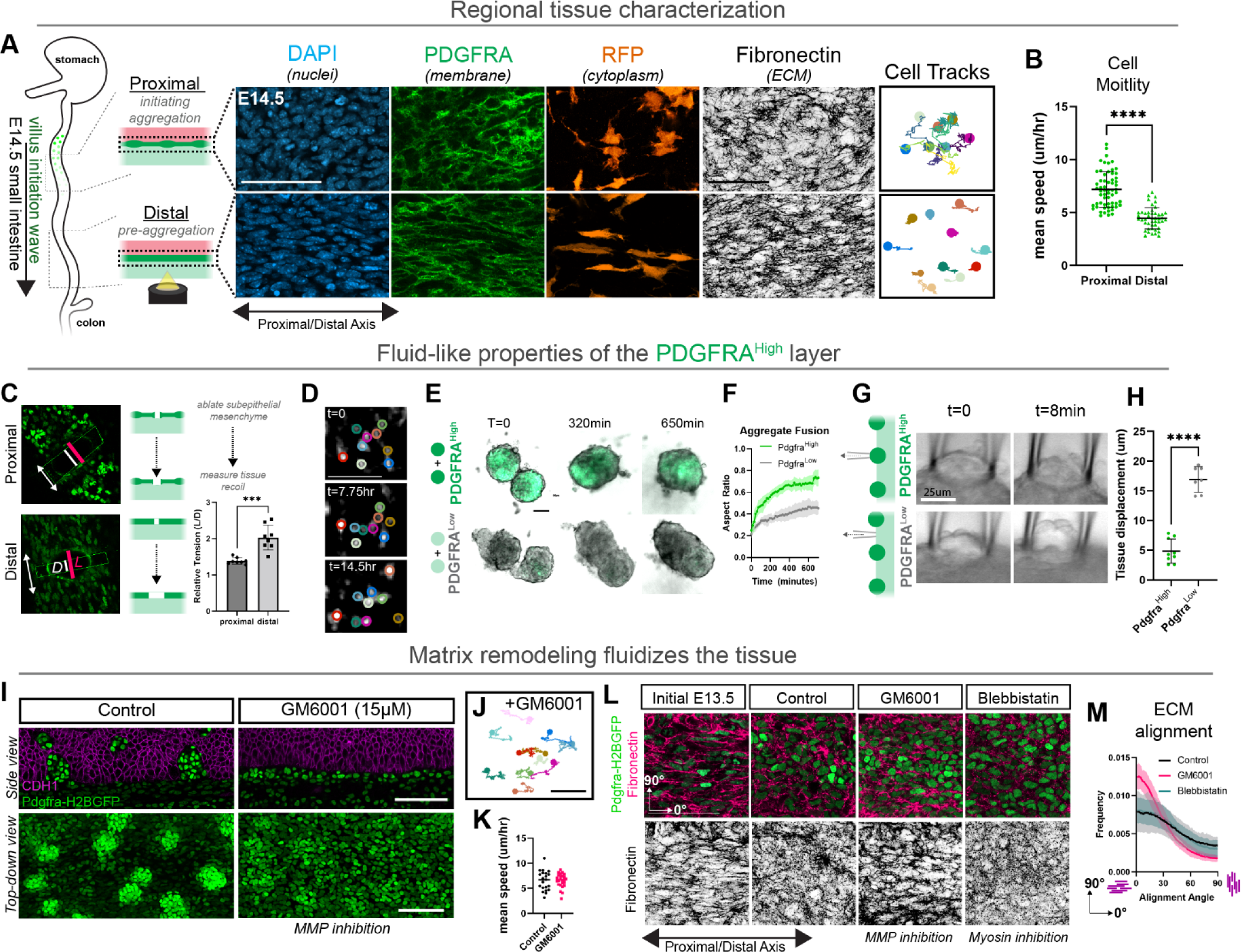
MMP activity fluidizes the subepithelial mesenchyme to enable cell aggregation and villus initiation through increased motility and the emergence of an altered surface tensions. (A) Spatial characterization of cell and ECM morphology and dynamics in the proximal and distal regions of the small intestine at E14.5. Cell motility tracks captured from region-specific timelapse movies over a 16 hour period. (B) Quantification of cell motility in proximal and distal subepithelial mesenchyme. (C) Quantification of relative tension as measured by the length of tissue recoil *L* (magenta line) at *t* = 60s of laser exposure relative to the initial distance *D* (white line) at *t* = 0s between the two edges of the maximum point (center) of the gap generated in the tissue by the laser cut, as shown by the yellow rectangle region of interest and white line. Optical sections taken immediately post laser ablation in proximal and distal subepithelial mesenchyme. Views are top-down *en face* views of the subepithelial layer. Double headed white arrow indicates the longitudinal (proximal-distal) axis of the gut. Each point represents a single ablation experiment done in a separate gut. (D) Neighbor exchange of PDGFRA^High^ subepithelial cells. Snapshots from a timelapse movie at noted times of the proximal intestine at E14.5. Each color represents a distinct cell. Note relative initial and final positions are different. (E) Coalescence of two PDGFRA^High^ spheroids over time. Snapshots are optical sections captured from a timelapse movie. (F) Quantification of aggregate fusion in panel E from timelapse movies. Data are represented as mean ± SD from *n =* 11 homotypic spheroid pairs used for each tissue type. (G) Widefield images of micropipette aspiration experiments. The PDGFRA^High^ tissue is less deformable than the PDGFRA^Low^ neighboring tissue. (H) Quantification of experiments in panel G. Each point represents a separate measurement performed on a separate gut fragment. Data are represented as mean ± SD (I) Optical sections from intestinal explants grown from E13.5 for 2 days in the presence or absence of MMP inhibitor GM6001. (J) Cell tracks from timelapse movies taken over a 16 hour period of E14 explants treated with GM6001. Compare to control tracks in (A). (K) Comparison of cell motility (mean speed) of cells in explants from C/D. (L) IF of optical sections (top) and single-channel maximum intensity projections of the first 10 µm of subepithelial mesenchyme (bottom) in freshly isolated or explanted tissues in noted treatment conditions grown from E13.5 for 2 days. (M) Quantification of Fibronectin (ECM) alignment relative to the proximal-distal axis of the gut (0 °) from experiment in (F). Data are represented as mean ± SD from *n = 9* measurements from distinct regions of interest in the proximal jejunum per condition from *n = 3* explants per treatment. Scale bars = 50 µm, *\*\*\*p<0.001; ****p<0.0001*

As cells, nuclei and ECM are known to align parallel to the direction of tensile stresses^36, 37^, these observed changes in nuclear and ECM anisotropy suggest a loss of residual stresses within the proximal intestine, consistent with a transition from an elastic to more fluid-like material regime among these two tissue regions. As would be expected in such a model, cell motility within these regions is markedly different. Specifically, cells within the distal non-aggregating gut display slower, confined motions in comparison to the proximal aggregate-forming tissue, where cells have faster, diffusive motility that extends beyond the diameter of a cell (**Figures 4A, 4B, and Videos S3 and S4**). We more directly tested for relative residual stresses in the proximal versus distal subepithelial mesenchyme at E14.5 by first removing the epithelium and then using a focused laser to generate precise cuts specifically within this subepithelial layer and without affecting the deeper tissues (e.g., smooth muscle). Distal and proximal tissues displayed different recoil behaviors: the distal tissue rapidly opened up to a greater degree along the proximal-distal axis compared with the proximal intestine (**Figure 4C**), consistent with the non-aggregating regions behaving more like an elastic solid that accumulates residual stresses under the influence of the extending gut axis^38, 39^. Thus, cell and matrix anisotropy are lost just prior to the onset of villus cluster formation, and this reduction in tissue alignment correlates with a decrease in residual stresses concomitant with an increase in cell motility.

### The PDGFRA^High^ tissue layer exhibits fluid-like properties and dewets from surrounding tissue

A thin layer of PDGFRA^High^ tissue separating from the surrounding PDGFRA^Low^ tissue into a patterned series of cell-ECM aggregates is qualitatively analogous to the dewetting of a thin film of liquid water on a non-wettable (i.e., hydrophobic) surface, where the high surface energy fluid film breaks up into a series of lower surface energy droplets^40^. In the case of the intestinal mesenchymal tissue, this appears to be an active process propelled by non-muscle myosin activity and the enhanced cohesive properties of the PDGFRA^High^ tissue. To directly test this concept, we examined whether the PDGFRA^High^ layer exhibits additional properties of a high surface energy fluid as would be necessary for active dewetting. In addition to the reduction of residual stresses as described above, we would expect **1)** diffusive (as opposed to concerted/collective) motility, **2)** merger and exchange of material within fluid-like tissue droplets, and **3)** elevated surface tension (itself driven by the high cohesivity of PDGFRA^High^ cells within).

We first examined whether PDGFRA^High^ cells undergo neighbor exchange (a hallmark of diffuse motility in fluid-like tissues)^41, 42^, as opposed to the correlated cell motions that would be expected for a contracting solid. By tracking individual cells at the onset of cell aggregation, we observed that PDGFRA^High^ cells tend to move non-collectively and swap neighbors on the timescale (<16 hours) of cell aggregation (**Figure 4D**). Further reinforcing this observation, cells continue to move after aggregating, exchanging positions between the interior and border of the aggregate (**Video S5**).

Second, we reasoned that two spherical aggregates of PDGFRA^High^ cells placed in contact would rapidly coalesce to form a single spherical aggregate, just as two droplets of fluid coalesce into a larger droplet. To test this, we microdissected PDGFRA^High^ cells away from surrounding tissues and grew them in suspension to generate spheroids containing PDGFRA^High^ cells and their native matrix but not neighboring cell types, such as the PDGFRA^Low^ cells and epithelium. The initially rough interfaces of these fragments rounded up to form spheroids (a process presumably driven by tissue surface tension and resisted by viscosity)^43, 44^, but occurred considerably faster in PDGFRA^High^ than PDGFRA^Low^ microtissues (**Figure S5B**). Subsequently, PDGFRA^High^ spheroids positioned in proximity quickly fused and coalesced to generate larger aggregates that also maintained a spherical geometry (**Figure 4E and 4F**). Coalescence occurred faster between PDGFRA^High^ spheroids than PDGFRA^Low^ spheroids, consistent with the two tissue types having distinct material properties. Further supporting a requirement for the ECM in the process of cell aggregation, as opposed to cell-cell interactions alone, any attempt at forming spheroids in suspension without native ECM failed.

Third, we tested whether PDGFRA^High^ aggregates had higher tissue surface tension than surrounding PDGFRA^Low^ tissue, which would be predicted if they behaved as a cohesive fluid. To do so, we performed micropipette aspiration on mesenchymal tissue fragments that had their epithelial layers removed such that we could access the mesenchyme and test its properties directly. We applied this method to both aggregating and non-aggregating tissue fragments and found that when applying the same pressure for the same duration, PDGFRA^High^ tissues deformed far less than the surrounding PDGFRA^Low^ tissues (**Figure 4G and 4H**). Taken together, these results support a model wherein the tissue undergoes a phase transition just prior to cell aggregation, from a solid-like state to a fluid-like state in a wave along the proximal-distal axis of the small intestine, enabling the PDGFRA^High^ mesenchyme to dewet from its neighboring interfaces.

### MMP-mediated ECM remodeling initiates fluidization of the subepithelial mesenchyme to enable cell aggregation and villus initiation

We hypothesized that this tissue phase transition is facilitated by the remodeling of ECM composition (from fibrillar to non-fibrillar, **Figure 2E**) and topology (from anisotropic to isotropic, **Figure 4A**), which would trigger the observed transition from confined and directional cell motility restricted to the axis of ECM alignment to a more rapid and less-biased (more diffusive-like) cell motility. Moreover, beyond this initial reduction of ECM alignment at the onset of aggregation, well-formed aggregates lack fibrillar ECM, further suggesting that ECM remodeling and a switch in ECM compositions could contribute to cell aggregation (**Figure S5C**).

Consistent with this idea, we observed an increase in expression of proteolytic genes and pathways during PDGFRA^High^ cell differentiation (**Figure S2M-P**). In particular, *Mmp2* and *Mmp14*, which encode matrix metalloproteinases that are known to degrade/remodel fibrillar ECM, are uniquely expressed by PDGFRA^High^ cells (**Figure 2E**). We therefore tested whether MMP activity is required for villus initiation by inhibiting MMP activity with GM6001 or Marimastat, two broad spectrum MMP inhibitors. MMP-inhibited explants failed to undergo mesenchymal aggregation and villus initiation (**Figures 4I and S5D**). Furthermore, MMP inhibition prevented the normally observed reduction in cell and matrix anisotropy during tissue maturation, and although cells remained motile, their movement was confined along the directionality of ECM fiber alignment (**Figure 4J-M**). Therefore, MMP activity drives villus initiation by promoting matrix remodeling and allowing for diffusive cell motility.

These results allow us to refine a model wherein the subepithelial tissue (comprising both cells and ECM) exists in two spatiotemporally distinct states and undergoes a phase transition from a more solid-like state to a more fluid-like one in a developmental wave along the proximal-distal axis of the intestine, triggering patterning and morphogenesis of the tissue interface. This fluidization event is driven by the activities of MMPs and actomyosin, which act alongside cell cohesion within the PDGFRA^High^ tissue to give rise to elevated tissue surface tension. Together, these emergent properties are sufficient to drive active dewetting of the PDGFRA^High^ tissue from PDGFRA^Low^ tissue in a process conceptually analogous to cell sorting^45–49^, but using different molecular machinery.

### A continuum framework for mesenchymal dewetting

The striking resemblance of PDGFRA^High^ cell aggregation to dewetting prompted us to seek more generalizable principles governing this process. Building upon our observations that this is an active, fluid-like tissue with elevated surface tensions between distinct tissue layers, we explored a continuum framework to formalize and test a conceptual dewetting model based on Cahn-Hilliard (C-H) phase separation dynamics. This class of model has been theorized to apply to tissue patterning^49, 50^ and has widely been used to describe dewetting in non-living systems^51, 52^, but has not been applied to directly reveal dewetting phenomena in tissue morphogenesis. In this model, PDGFRA^High^, PDGFRA^Low^, and epithelial tissue types are represented by distinct material phases that reorganize based on diffusive motion within the material, directed by differential surface tensions arising from cohesion and adhesion^46, 53^ (**Figure 5A**). Thus, we parameterized interfacial tensions PDGFRA^Low^-PDGFRA^Low^ > PDGFRA^High^ -Epithelium > PDGFRA^High^-PDGFRA^Low^ (where higher tension represents a less stable interface, see **Supplementary Text** for details) based on adhesion molecule expression and the biophysical measurements of surface tensions and cohesivity described above. We initialize the model using the tissue configuration seen *in vivo* before the onset of morphogenesis, wherein the PDGFRA^High^ layer is sandwiched between the epithelial and PDGFRA^Low^ layers. We then introduce noise into the PDGFRA^High^ phase (i.e., tissue compartment), inducing its breakup into distinct droplets with stereotyped shapes that produce curvature at the surrounding interfaces, recapitulating tissue morphogenesis (**Figure 5A and 5B**). *In vivo*, this initiating noise is interpreted to be non-muscle myosin-based activity that manifests as non-directed, diffusive-like cell motility, which at the continuum limit becomes equivalent to the active diffusion of particles. Indeed, without noise (as without myosin activity), droplets fail to form (**Figure S6A)**. Remarkably, the breakup into clusters occurs robustly as long as the surface energies are favorable, a condition that is satisfied by a very broad range of surface energies (**Figure 5A, S6A and S6B**). As would be expected from a dewetting mechanism, PDGFRA^High^ aggregates within the model and *in vivo* evolve characteristic geometries and contact angles at the tissue interfaces during aggregate maturation, and these values shift when the interfacial properties are modified **(Figure S6C-G)**.

**Figure 5.**
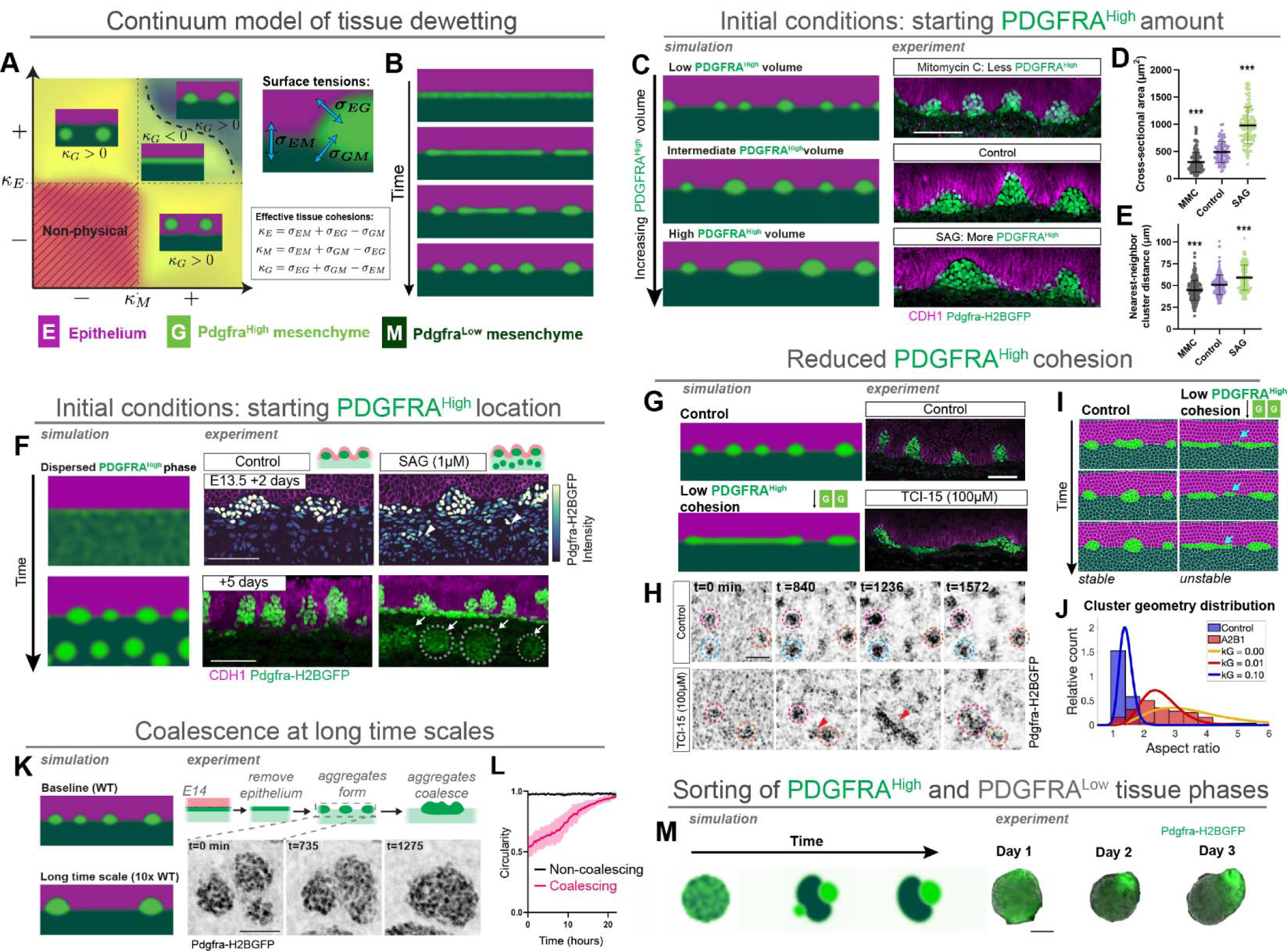
Computation modeling predicts outcomes of cell aggregation and interfacial morphogenesis through mesenchymal dewetting. (A) Stable clustering can occur if and only if the effective cohesion is positive for all three phases. If one of the cohesions is negative, clustering may occur, but the clusters detach from the interface. If two or more cohesions are negative, the system is non-physical and unsolvable. *Κ* denotes the effective tissue cohesion values for each tissue layer. (B) Breakup of the initial noisy PDGFRA^High^ layer into distinct clusters in the wild-type simulation. (C) Simulation and experiment demonstrating the effect of initial PDGFRA^High^ volume on the outcome of cell aggregation. To minimize the effects of coarsening, the simulations are run until the time point when the clusters first become clearly separated and assume a close-to-final droplet-like shape, considered the wild-type simulation as demonstrated in panel B. Explants were treated with noted compounds to increase or decrease the amount of PDGFRA^High^ cells at the onset of aggregation. Images are representative of *n =* 6 tissues per condition. See detailed explanation in Methods. (D) Quantification of aggregate size calculated by cross-sectional area at the midplane (maximum) of the aggregate in 3D whole mount tissues. Each dot represents a single cluster, measured from 4 tissues per condition. Data are represented as mean ± SD (E) Quantification of cluster spacing by nearest neighbor detection, measured from 4 tissues per condition. Data are represented as mean ± SD (F) Initializing PDGFRA^High^ material deep into the mesenchymal phase in the simulation results in the formation of deep ectopic clusters below the normal interfacial clusters. Treating the tissue with SAG over the course of 5 days similarly results in formation of deep ectopic clusters, noted by white arrows and dashed lines to indicate sphericity. (G) Continuum (C-H) simulation of lowered PDGFRA^High^-PDGFRA^High^ cohesion and experiment of explanted tissues treated with TCI-15 from E13.5 for 3 days in culture. (H) Individual snapshots from a timelapse video highlighting diffusive cluster behavior in explant tissues treated with TCI-15, as opposed to stationary clusters in controls. (I) Time resolved cell-based simulations of lowered PDGFRA^High^-PDGFRA^High^ cohesion highlighting diffusive clusters that generate gap events between clusters, similar to those seen *in vivo* in (H). Blue arrows highlight gap events, consisting of breakup and fusion of cells from aggregates as they continuously model. No gap events are present in the wild-type model where aggregates are stable. (J) Quantification of aggregate aspect ratio *in silico* and *in vivo*, measured at the aggregate midplane of maximum epithelial deformation in whole-mount tissues. Bars of the histogram are representative of the control and experimental data from Figure 3H, and the lines represent inverse Gaussian fits to the cell-based simulations of respective cohesion energies, demonstrating close agreement between predicted and measured geometries. (K) Long-timescale coalescence of the clusters in the model, and the coalescence of clusters in the tissue when grown without epithelium. (L) Quantification of aggregate geometry during coalescence, measured from the onset of two or three aggregates coming into contact and compared with non-coalescing clusters. Control (non-coalescing) clusters were measured from those aggregates that did not contact and fuse with other clusters during the duration of the videos. Data are represented as mean ± SD; *n = 12* coalescing clusters and *n = 6* non-coalescing clusters. (M) Simulation of phase separation in a reduced two-phase system consisting of initially mixed population of high-cohesion PDGFRA^High^ material and lower cohesion PDGFRA^Low^ material, and the corresponding experiment with microtissue spheroids initiating with a mixed population of PDGFRA^High^ and PDGFRA^Low^ cells. Experimental data is representative of *n = 11* mixed spheroids. Scale bars = 50 µm, *\*\*\*p<0.001*

### The properties of an active and cohesive mesenchymal fluid patterns and shapes the tissue interface

We used the C-H model to further explore the properties of mesenchymal dewetting and the impact of altering tissue-level parameters on villus formation. To begin, we modulated the initial volume of the PDGFRA^High^ layer, finding that this impacts aggregate size and spacing. Specifically, a lower input volume results in smaller aggregates that are spaced closer together while a higher input volume generates larger aggregates spaced farther apart (**Figure 5C**). To test these predictions in living tissues, we treated intestinal explants with compounds that alter the amount of aggregate-forming PDGFRA^High^ cells at the tissue interface. We took advantage of our previous observation that blebbistatin treatment uncouples PDGFRA^High^ cell differentiation from tissue morphogenesis in a reversible manner (**Figure 3D and S4C**). This allowed us to synchronize tissues such that, upon blebbistatin washout, we could observe the direct effect on pattern formation of having either fewer or more PDGFRA^High^ cells (see detailed explanation in Methods). To reduce the amount of PDGFRA^High^ cells at the subepithelial interface, we exposed tissues to a brief pulse of mitomycin C (MMC), an irreversible cell cycle inhibitor (**Figure S6H)**, resulting in aggregates that were smaller and spaced closer together than in control tissues (**Figure 5C-E**). To generate a greater amount of PDGFRA^High^ cells, we treated tissues with either SAG or Dorsomorphin (a type I BMP receptor inhibitor), treatments known to generate larger cell aggregates^9, 19^. Consistent with computational modeling, we additionally found that these treatments resulted in a higher amount of mesenchymal cells at the interface that subsequently broke up into aggregates that were larger and spaced farther apart (**Figure 5C-E and S6H-L**). Therefore, the pattern of cell aggregation, and thus the initial field of emerging villi, can be specified directly as a function of the number of PDGFRA^High^ cells present at the initiation of tissue fluidization, without the need to invoke a chemical-based reaction-diffusion patterning mechanism.

We then tested the importance of the localization of PDGFRA^High^ cells at the onset of morphogenesis. These cells normally differentiate as a thin band adjacent to the interface with the epithelium (**Figure 5A**). We found that when the model is initialized with a PDGFRA^High^ phase that instead extends deeper and is mixed within the normally PDGFRA^Low^ phase, the outcome of morphogenesis is altered. Rather than a regular pattern of aggregates at the interface, circular ectopic aggregates of the PDGFRA^High^ phase form deeper within the tissue (**Figure 5F**). Recognizing that mesenchymal cells deeper in the tissue are also Hh responsive^19, 27, 54^ (**Figure S3H, S3L, and S3M**), we reasoned that longer incubations with SAG would lead to ectopic differentiation of PDGFRA^High^ cells. Consistent with this notion and adding further support for the fluid-like behavior of these cells, PDGFRA^High^ cells were induced by SAG treatment deeper within the mesenchyme and eventually formed ectopic clusters with characteristic droplet-like spherical geometries in the deep tissue (**Figure 5F, S6E and S6F**). Therefore, Hedgehog activity levels, which are normally highest in the mesenchyme adjacent to its source in the epithelium^19, 27^, are sufficient to activate the PDGFRA^High^ program at saturating concentrations throughout the mesenchyme, and subsequently these ectopic cells round-up to form spherical aggregates as they separate from the PDGFRA^Low^ tissue phase.

Next, we asked whether the high cohesion of PDGFRA^High^ cells is necessary for breakup of the subepithelial monolayer, aggregate formation, and epithelial deformation. The model predicts that with reduced PDGFRA^High^ monolayer cohesion, breakup still occurs, but the aggregates fail to adopt a regular geometry, compact, and generate interfacial folds (**Figure 5G**). The results of these simulations bear a striking resemblance to those of tissues treated with the A2B1 Integrin inhibitor TCI-15, which blocked aggregate compaction and interface folding and led to wider and more heterogeneous cell aggregates (**Figure 3F-H and 5G**). By live imaging explants, we found that in addition to impairing cluster geometry, TCI-15 treatment prevented the tissue from reaching a stable steady-state: the emerging clusters continuously remodeled, with individual cells moving between clusters and clusters fusing and breaking up, whereas aggregates are highly stable once formed in control tissues (**Figure 5H and Video S6**). These fluctuations manifest in the movements of discrete units of cellular mass, in contrast with the C-H model, which describes the material flow in continuum units of mass manifesting at the tissue-level. Therefore, to closer investigate this continuous remodeling observed at the cell-level, we simulated the activity of individual cells and the resulting dynamics in a cell-based approximation of our continuum model, using a self-propelled Voronoi model^41^ (SPV; see **Supplementary Text**). As with the C-H model, the SPV model breaks up into a series of stable aggregates at the interface between the cell layers (**Figure S6M**), generates interfacial curvature, and furthermore recapitulates the above predictions of aggregate patterning (**Figures S6N-Q**). In control SPV simulations, aggregate formation is highly stable and cells do not move between aggregates; however, when PDGFRA^High^ cohesion is lowered, the simulated aggregates continuously exchange cellular material among neighbors (**Figure 5I**), matching the cell dynamics and aggregate geometries observed by timelapse microscopy in explant tissues treated with TCI-15 (continuous remodeling of aggregates) versus controls (stable aggregation) (**Figure 5H and 5J**). The close agreement between the structure and dynamics of clusters further supports a conceptual model in which high PDGFRA^High^ cohesion, mediated by integrins, is critical for generating stable aggregates and the surface tensions necessary to fold the overlying interface. However, high cohesion is dispensable for breakup of the PDGFRA^High^ tissue layer.

The C-H model predicts that the PDGFRA^High^ phase coarsens over long timescales to eventually resolve into fewer, yet larger clusters, through energy minimization – just as would be expected through the fusion of fluid droplets during dewetting (**Figure 5K**). This contrasts with *in vivo*, where the aggregate and villus positions, once established, are largely fixed (**Figure 5H**). We reasoned that the nature of this difference was due to the epithelial phase also being modeled as a fluid within the C-H model, whereas *in vivo* the epithelium is largely non-motile and therefore might function to restrict the diffusion of the PDGFRA^High^ clusters (**Figure S2A**). We tested this observation experimentally by returning to our finding that the PDGFRA^High^ mesenchyme forms aggregates at the cell-media (i.e., liquid) interface after removal of the epithelium (**Figure 1J**), a condition that mimics the lack of a low activity overlying tissue layer that we suspected could resist aggregate movement. Strikingly, when epithelial-free mesenchyme is grown for several days following condensation, the aggregates remained motile, contacted one another, and coalesced to form larger circular aggregates where cells comingled (**Figure 5K, 5L and Video S7**), reinforcing our previous observations using microtissue spheroids (**Figure 4E**). Confirming these cell-level phenotypes *in silico*, enabling motility of (i.e., fluidizing) the epithelium within the SPV model also results in aggregate coalescence (see **Supplementary Text and Figure S6R**). Thus, the epithelium serves as a mechanical anchor for the clusters: in order for clusters to move in the presence of epithelia, activity (impulses) generated by PDGFRA^High^ aggregates must be sufficient to “push” the overlying epithelium out of the way – a condition that is relaxed when we fluidize the epithelium (models) or remove it altogether (*in vivo*).

A central element of the proposed active dewetting mechanism is that the driving forces at the tissue scale are conceptually analogous to those driving cell sorting, whereby two initially mixed populations separate into discrete populations based on their interfacial properties^45, 46, 49^. Indeed, we can recapitulate cell sorting using the C-H model by reducing it to a two-phase system comprising only PDGFRA^High^ and PDGFRA^Low^ phases. Starting from initially well-mixed populations as in models of cell sorting, the model evolves through phase separation to generate a PDGFRA^High^ pole at one edge of the PDGFRA^Low^ phase, sometimes passing through an intermediate stage with multiple PDGFRA^High^ poles that eventually coalesce. We tested this experimentally by generating spheroids as in **Figure 4E** but from earlier tissue that had yet to aggregate. These spheroids contained two evenly distributed cell types and evolved in time in a manner qualitatively similar to the model (**Figure 5M**). In addition, the more rapid coalescence of PDGFRA^High^ homotypic spheroids (**Figure 4E**) is accurately predicted by the two phase model (**Figures S6S and S6T**). Taken together, the close agreement between modeling and experiments supports that PDGFRA^High^ cells behave like a cohesive and active fluid that dewets from its surrounding interfaces into a series of aggregates. The high cohesion results in an elevated surface tension of aggregates that then promotes their rounding, while the coupling of their surface tension with the overlying epithelial layer initiates folding at the tissue interface, marking the position of future villus outgrowth.

## DISCUSSION

Tissue folding at epithelial-mesenchymal interfaces is widespread throughout organ development. Folds often emerge in patterns that are explained as resulting from a Turing-like reaction-diffusion mechanism of signaling molecules or mechanical instabilities^1, 8, 49, 55–58^. A fundamental question in developmental and regenerative biology is how these two processes – establishing the positional information and the geometry – can occur robustly across scales and within diverse signaling and mechanical environments. There are likely many ways to fold a tissue, and understanding these modes can lay a foundation for bioengineering strategies toward building functional tissue interfaces or reactivating these programs following tissue damage. Here, we identify a mechanism involving the active and fluid-like properties of a thin layer of mesenchymal cells that is sufficient to program both pattern and geometry of the epithelial-mesenchymal interface in the developing intestine to initiate the formation of villi.

### Forces and signals shape intestinal villi

Distinct mechanisms have been hypothesized to underly villus folding in mammals versus birds^18^. In the chick, highly aligned smooth muscle layers constrain the faster growing inner endoderm and mesenchyme, causing them to buckle into a series of folds^8^. While this same framework was proposed to account for the emergence of villi in the mouse intestine^8^, more recent data, including those presented here, indicate that this model fails to account for the initial folding event^12^. These findings led us to investigate an active role for mesenchymal PDGFRA^High^ cell aggregates, termed villus clusters, that form at the tissue interface at the same time and place where the epithelium begins to fold into villi^10, 19, 59^.

Villus clusters have been proposed to act as signaling centers to activate shape changes in the overlying epithelium through currently unidentified signals^12, 22, 60^. In contrast, we find that the initial interfacial folding event can be explained by the emergent interfacial properties of cohesive PDGFRA^High^ mesenchyme. Specifically, the emergence of an elevated surface tension in this fluid-like tissue provides a driving force for a reduction in its surface-to-volume ratio, leading to aggregate formation. Simultaneously, the overlying interface is deformed through aggregate surface tension to generate stereotyped patterns and geometries of curvature. Our data suggest that this initial epithelial deformation effectively traps the aggregates locally in a process that we suspect may be further reinforced by mechanical changes in the epithelium, while the local patterning of proliferative zones induced by tissue curvature likely helps maintain villus cluster pattern and villus elongation^13, 60, 61^. In support of this, we observe the apical deformation that occurs in normal villus outgrowth is not present when the mesenchyme is recombined with the colonic epithelium, further suggesting that unique epithelial properties underly differences between colonic (which lacks villi) and intestinal luminal topographies *in vivo*.

In addition to establishing a mechanical framework for villus initiation, our work provides insight into how the underlying cell behaviors are genetically programmed to occur with spatiotemporal precision. Further, our work adds mechanistic context to signals previously implicated in the process. One such signal is Hedgehog (Hh), which has a demonstrated, yet not clearly defined, involvement in villus initiation^19, 54, 62, 63^. Here, we show that Hh is sufficient to induce PDGFRA^High^ cell differentiation and mesenchymal aggregation (at both the tissue interface as well as deep within the mesenchyme) in the absence of any other epithelial cues. How Hh exerts these effects is not clear but it may act through downstream Planar Cell Polarity (PCP) signaling^22, 54^. To this end, the transcriptional dataset collected here, that captures the entirety of this timeframe including all major cell types of the intestine, will enable future examination of these interactions.

Villi form in sequential rounds^19, 61^, and we hypothesize that these subsequent events result from the exposure of deeper mesenchymal cells to sufficiently high levels of inductive Hh signaling as the inter-villus epithelium pushes into the mesenchyme. This view is in contrast to previously invoked Turing-like reaction-diffusion mechanisms, in which BMP pathway ligands and antagonists emanating from the forming aggregates have been proposed to generate the observed spot pattern of clusters^9^. Instead, we find that aggregate pattern (i.e., spacing and size) is set by the biophysical properties of the system as well as the initial conditions – namely the amount of PDGFRA^High^ cells at the onset of mesenchymal aggregation – and further show that both BMP inhibition and Hh activation modulate this parameter.

### Dewetting and phase transitions in tissue biology

Drawing analogies between the behaviors of living tissues undergoing shape and size changes and the physics of non-living systems can provide new insights into the mechanisms that drive morphological changes. Here, we draw upon the analogy of a thin liquid-like film that dewets into droplets to describe how a thin layer of PDGFRA^High^ cells at the epithelial-mesenchymal interface transforms into a series of droplet-like aggregates, just as rain on a leaf or windshield forms distinct droplets - processes driven by the minimization of surface energies. Wetting behaviors and transitions are likely widespread in developing tissues where they apply to the spreading (wetting) or retraction (dewetting) of cells on substrates of varying properties^64–68^.

A precondition for these behaviors is that the tissue can act in a fluid-like manner. Accordingly, we find that the intestinal mesenchyme undergoes a phase transition from a solid-like state in the immature, distal tissue, to a fluid-like state in the more mature proximal tissue, at the precise time when the cells begin to aggregate. Such tissue phase transitions enable spatially graded changes in material properties that give rise to unique morphogenetic events, but these have so far been primarily studied in epithelia or dense mesenchyme lacking ECM^69–76^. In contrast, the subepithelial mesenchyme of the gut undergoes an unusual and unique phase transition, as it requires a composite of cells and ECM to remain fluid throughout aggregation. Moreover, instead of relying on previously described mechanism of tissue fluidization, such as cell density or cell-contact fluctuations, fluidity is initiated within the gut mesenchyme through the function of locally expressed matrix remodeling factors. How this phase transition is coordinated in a developmental wave across the proximal-distal gut axis, as well as how the cell-ECM composite remains fluid, is an exciting area for further investigation that we speculate involves a switch from fibrous to non-fibrous ECM. In support of this mechanism, we observe a transcriptional shift in genes encoding these ECM types, including the downregulation of fibrillar collagens and the upregulation of *Col23a1*, which encodes a transmembrane collagen that binds ITGA2B1 and regulates cell cohesion in the skin^77^.

Given that the tissue undergoes a solid-to-fluid transition to initiate villus formation, it is reasonable to believe that the tissue can go through a fluid-to-solid transition later in development to maintain and progressively rigidify villus form, a question that we have not completely addressed here. Interestingly, however, across the timescales of villus initiation (E14.5-E16.5), we observe that mesenchymal aggregates remain fluid-like as opposed to solidifying. Thus, active dewetting results in a pseudo-steady state that will continue to evolve in time towards larger aggregates unless locked into place by subsequent morphogenetic events. This is exemplified by the observation that aggregates coarsen through coalescence over time toward a lower surface-to-volume ratio in the absence of a constraining epithelium. Active and/or passive interfacial properties are likely important for the final morphologies of folds in other systems as well. For example, fine-tuning Wnt signaling levels in the developing tooth determine whether the tissue evaginates or invaginates at the site of condensates^78^, and in the mammalian skin, reducing epithelial contractility results in a failure of epidermal hair placode invagination and leads to folds above dermal condensates that are strikingly similar to those in the intestine^79^. Thus, by altering the passive or active properties of neighboring tissue layers, as well as those of the aggregating cells, tissues can self-organize into a diversity of folded geometries and patterns. Indeed, in our previous work we have highlighted the versatility of mesenchymal compaction as a mode to generate both physiologically relevant and highly stylized tissue patterns^80^.

### Mesenchymal condensations mediate tissue morphogenesis

Mesenchymal condensations at tissue interfaces are widespread throughout organogenesis, including in the developing kidney^81^, lung^82^, tooth^83^, musculoskeletal tissues^84^, and skin^2, 85^, where they play multiple roles in determining the location, rate, and geometry of forming tissues. Classical tissue recombination experiments have highlighted the inductive capacity of mesenchymal condensates to drive tissue identity in overlying epithelia^86^, and reconstitution experiments have demonstrated the remarkable ability of such condensates to serve as organ precursors *in vitro*^87, 88^. Understanding the cellular mechanisms underlying mesenchymal condensations across diverse contexts *in vivo* will inform *in vitro* engineering strategies aimed at generating true-to-form and complex organs and organoids with functional tissue interfaces.

Cellular condensation has classically been thought to result from epithelial pre-patterns that signal to the underlying mesenchyme to organize. However, we find that condensates in the embryonic intestine can initiate spontaneously and autonomously within a self-organizing mesenchyme, while signals (e.g., Hh) from the epithelium specify the positional domain within the mesenchyme permissive for aggregation. While we demonstrate that the initial patterning and folding of the tissue interface can occur through this mechanism independently of a genetic prepattern or chemoattractive cues, such as those invoked in other condensate-forming contexts^83, 89, 90^, we cannot rule out the possibility that pattern reinforcement or the pace of aggregation, once nucleated by mesenchymal dewetting, could involve such chemical cues, possibly as secondary stabilizing mechanisms.

Strikingly similar mesenchymal condensation events are observed during the formation of dermal papillae, periodically patterned mesenchymal aggregates involved in the formation of hair follicles that bear morphological resemblance to villus cluster condensates^10^. As in the case of villus clusters, dermal papilla formation coincides with mesenchymal cell motility, subsequent exit from the cell cycle, and can occur in the absence of an apparent epidermal prepattern, suggesting that mesenchymal condensation here might also occur autonomously^89–92^. This apparent self-organizing ability of the dermis has been studied with more detail in the chick skin, where dermal condensates initiate feather bud morphogenesis^85, 93–95^. While the overall morphological outcome – mesenchymal condensation – is qualitatively similar to the aggregation of PDGFRA^High^ cells of the gut, the underlying mechanisms appear distinct. Specifically, dermal condensate formation relies on supracellular contractility and alignment of ECM to drive the collective flow of cells and ECM into the aggregate^95^. In contrast, mesenchymal cell motility in the gut is enabled by the loss (rather than gain) of ECM alignment; cells move individually in a non-coordinated manner (rather than as a supracellular collective); and the resulting aggregates have reduced (rather than concentrated) ECM. Furthermore, while models for mesenchymal condensation of both the avian dermis and the mammalian gut invoke fluid-like tissue behaviors, highlighting the generalizability of this framework, the gut tissue is unique in that it both patterns and folds due to the macroscopic manifestation of cells interacting through high cell-cell cohesions — the emergent surface tensions. This active dewetting mechanism is therefore additionally remarkable because it operates at tissue interfaces rather than in driving density heterogeneities within the bulk tissue, enabling the simultaneous sculpting and patterning of such interfaces into diverse, functional forms.

## METHODS

### Mouse genetics

All animal experiments were performed in accordance with the guidance established by the Institutional Animal Care and Use Committee (IACUC) and Laboratory Animal Resource Center (LARC) of the University of California, San Francisco, CA, USA. For generation of embryos, the observed plug date was considered embryonic day 0.5. Tamoxifen (0.15 mg/g of body weight) was administered by oral gavage to pregnant dams consecutively at E9.5 and E10.5 to induce recombination in experiments using tamoxifen-inducible Cre lines. All strains were maintained on a C57BL/6J (Jackson Laboratory: 000664) background. Mice strains used in this study are listed below:

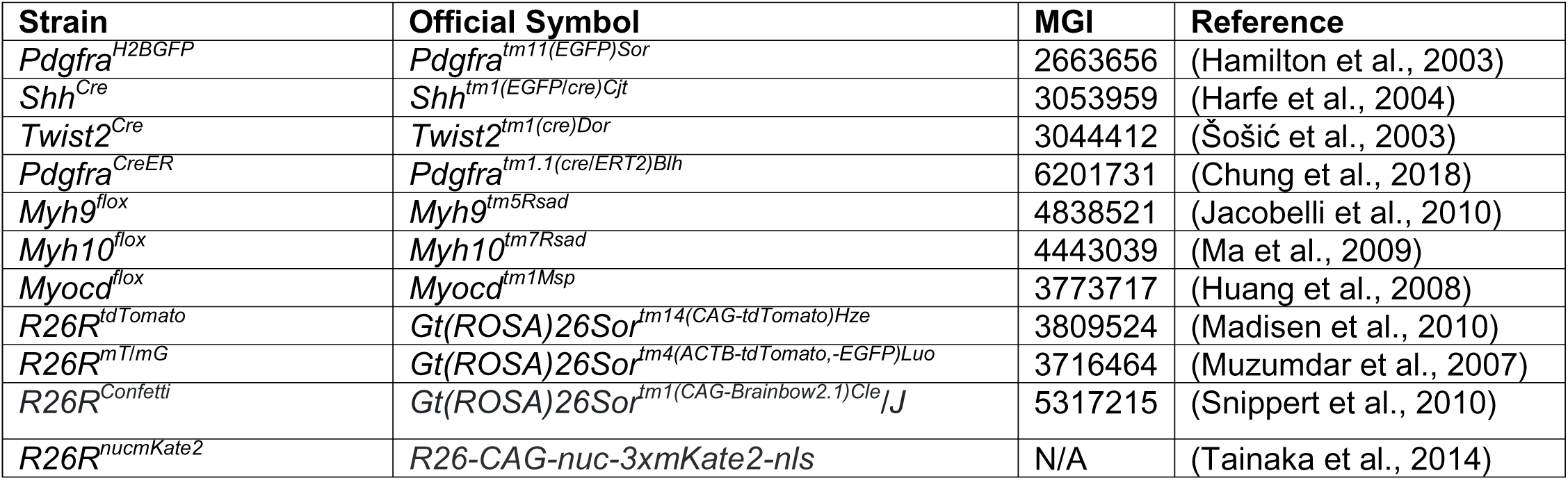
Summary of mouse alleles.

### *Ex vivo* explant tissue cultures

For standard explant cultures, embryonic intestines were harvested at specified timepoints and grown on transwell inserts at the air-liquid interface as described with slight modifications^96^. Specifically, isolated intestines were placed on 8.0 um pore-size polycarbonate transwell inserts (Corning) in 6 well plates containing 1.5mL fetal gut media (BGJb media containing 0.1 mg/mL ascorbic acid) and cultured for specified durations at 37C with 5% CO_2_. Media was changed twice a day, approximately every 12 hours, with fresh small molecules added at each media change. The following small molecules were used at the specified concentrations, unless noted otherwise in the figure, and control tissues were cultured in equivalent concentrations of vehicle (DMSO or water): SAG (Tocris, 1µM); blebbistatin (Sigma, 5µg/mL); Y27632 (Tocris, 20µM); para-amino-blebbistatin (Cayman Chemicals, 25µM); TCI-15 (100µM); GM6001 (Tocris, 15µM); Marimastat (Tocris, 15µM); Mitomycin C (Tocris; 3µg/mL); Aphidicolin (Cell Signaling Technologies; 10µM); Dorsomorphin dihydrochloride (Tocris, 12.5µM). To monitor the development of tissue and to ensure that cell clustering had not already initiated, dissected tissues were visualized by epifluorescence prior to the experimental start. For standard explant cultures, tissues were dissected at E13.5 and cultured for 48 hours, unless otherwise noted.

To synchronize morphogenesis in tissues in Figure 5C so as to specifically modulate the amount of starting material (i.e., PDGFRA^High^ cells) at the initiation of clustering, whole explanted intestines were cultured in the presence of 5µg/mL blebbistatin for two days. This treatment successfully blocks morphogenesis, yet allows for PDGFRA^High^ cells to differentiate in a thin layer at the interface of the mesenchyme and epithelium (see **Figure S6H**). Tissues were concurrently treated during this two day window with compounds detailed below to modulate the amount of PDGFRA^High^ cells at the interface. Following two days of incubation, a single layer of cells expressing high levels of PDGFRA was visible adjacent to the epithelium, but no deformation had occurred. To initiate morphogenesis, blebbistatin/compound-containing media was removed and replaced with basal fetal gut media and washed 6 times for 10 minutes each on an orbital shaker within the incubator at 100rpm. Following these washes, the tissue was incubated for another 24 hours in basal fetal gut media in the absence of any compounds. During this time frame, the mesenchymal cells aggregated and deformed the overlying epithelium. To inhibit proliferation, and thereby decrease the amount of PDGFRA^High^ cells as starting material, freshly isolated tissues were immediately incubated in fetal gut media containing 3µg/mL of the irreversible cell cycle inhibitor Mitoymcin C (MMC) for 1 hour in suspension on an orbital shaker within an incubator at 37C and 5% CO_2_. Subsequently, tissues were washed 6 times for 10 minutes each in the same orbital shaker setup while in suspension to remove residual Mitomycin C, which we found to be toxic to the tissue if not completely removed. Following these washes, the MMC-treated tissues were transferred to transwells and cultured as above with blebbistatin and subsequent washout. To increase the amount of starting PDGFRA^High^ cells at the interface, tissues were treated with either 1µM SAG or 12.5µM Dorsomorphin in combination with the blebbistatin treatment, and these compounds were also subsequently washout out (along with belbbistatin) after 48 hours of incubation.

### Live imaging

*Pdgfra^H2BGFP^; Shh^Cre^; R26R^tdTomato^* mice were used to visualize mesenchymal cell clusters and the epithelial layer, labeled by *Shh^Cre^*, simultaneously. For tracking of single nuclei, *Pdgfra^H2BGFP^*mice were used to label the mesenchyme, *R26R^nucmKate^*^2^ to labeling the epithelium, and for tracking of cells, *Pdgfra^H2BGFP^; Twist2^Cre^; R26R^Confetti^* mice were used to sparsely label mesenchymal cells with cytoplasmic RFP. *Twist2^Cre^* leads to recombination broadly throughout the intestinal mesenchyme and is apparent in nearly early *Pdgfra^H2BGFP^*-expressing cells, thus enabling the co-labeling of nuclei and cytoplasm of PDGFRA-expressing cells with different colors to enable tracking. The cytoplasmic YFP signal is also visible from the confetti reporter, but is dim compared to the *Pdgfra^H2BGFP^* signal. Notably, *Twist2^Cre^* also labels other PDGFRA-cells deeper within the mesenchyme, so care was taken to specifically image only the top layers of the subepithelial mesenchyme, which is composed nearly entirely of PDGFRA^High^ and PDGFRA^Low^ cells. Imaging was performed on E14 (no visible clusters at experiment start) embryonic intestinal explants that had been longitudinally cut open using tungsten needles (0.004in diameter, A-M systems). The fileted intestinal explants were cultured on Corning 8.0µm pore-size polycarbonate transwell membrane inserts in 6 well plates containing 1.5mL fetal gut media (BGJb media containing 1% Pen/Strep and 0.1 mg/mL ascorbic acid with the epithelial surface facing up). For imaging of aggregate fusion events in epithelial free conditions, timelapse movies were initiated after the mesenchyme-only tissues had matured for 2 days in culture such that the aggregates were already present at the start of the timelapse. 70µM of nifedipine was added to the culture media in all conditions to inhibit spontaneous smooth muscle contractions and therefore minimize tissue movement to enhance stability. Notably, we find that impairing smooth muscle inhibition with nifedipine does not impair cell differentiation and tissue morphogenesis (see Figure S1). To block non-muscle myosin activity in specific experiments, tissues were treated with 25µM para-amino-Blebbistatin, a more stable and less-phototoxic analog of blebbistatin (Cayman Chemicals), or 15µM Y27632, which indirectly inhibits non-muscle myosin activity. For epithelial-free experiments, 1µM SAG was added to the cultures to maintain mesenchymal viability. Images were acquired using a 4x(0.4NA) AZ Plan Apo Air long working distance objective with both optical and digital zooms on a Nikon AZ100M upright C2 confocal within an environmental chamber maintained at 37C with 5% CO_2_. Tissues were imaged for up to 48 hours depending on the experiment and were imaged at fixed intervals (either 12, 15 or 20 minutes depending on the experiment).

### Analysis and quantification of live imaging data

For the analysis of epithelial curvature and aggregate surface-to-area ratio, live imaging data obtained with *Shh^Cre^, Pdgfra^H2BGFP^, R26R^tdTomato^* reporter mice tissue were used. Regions of interest at edges of the tissue, where the midline of the cluster could be captured in optical sections as in Figure 1B to enable for visualization of both epithelial and mesenchymal layers in a single optical plane were used for quantification. Five regions of interest, representing single optical sections through five different aggregates with the clearest epithelial-mesenchymal boundary were used for the analysis. To measure the curvature of the epithelium, the epithelial-mesenchymal interface boundary immediately above aggregating PDGFRA^High^ cells was traced manually using ImageJ/Fiji. The resulting region of interest (ROI) therefore consisted of a region of curvature that was then fit to a circle, and the radius of curvature of the circle was calculated. The inverse of this value was plotted in Figure 1C as “Epithelial Curvature”. To measure the perimeter (i.e., surface) -to-area ratio, the same images were converted to binary and thresholded to specifically isolate PDGFRA^High^ cells. After applying a Gaussian blur, the total perimeter and area of thresholded pixels within a 60×60µm area of the mesenchyme at the sites directly beneath the epithelium containing each resulting aggregate were calculated for each captured frame, at 1 hour intervals, of the timelapse and plotted in Figure 1C.

For cell tracking, timelapse movies were first corrected for tissue drift and movement using StackReg plugin^97^ in ImageJ/Fiji. The cell nuclei (*Pdgfra^H2BGFP^*) or mesenchymal cells with mosaically labeled cytoplasms (*Twist2^Cre^; R26R^Confetti^*) were then tracked using semiautomated manual tracking in TrackMate^98^. Mean speed, total displacement, and *Pdgfra^H2BGFP^* intensity was calculated for individual nuclei in TrackMate across a 20 hour time period captured every 12 minutes.

To measure aggregate coalescence in the absence of the epithelium, aggregate circularity (4pi(area/perimeter^2) was measured in ImageJ/Fiji by first thresholding on the *Pdgfra^H2BGFP^* signal from timelapse movies. Subsequently, a median filter was applied to the thresholded images, and circularity was measured across the span of the movie for fusing aggregates composed of an initial two or three aggregates. Fusion events of more than three initial aggregates were excluded from quantification as these tended to reach circularity at longer time frames that extended beyond the duration of the timelapse videos.

### Epithelial-mesenchymal tissue recombination experiments

E13.5 small intestinal tissues were harvested in Calcium/Magnesium free (CMF) PBS and washed several times in CMF PBS before being placed in 3mL of Cultrex Organoid Harvesting Solution (R&D Systems), a non-enzymatic basement membrane depolymerization solution, within a 35mm dish. Tissues were subsequently incubated at 4C for 30-45 minutes (we note lot-to-lot variability in effectiveness) on an orbital shaker. Following incubation, tissues were kept on ice and filleted open using a 0.004in or 0.002in diameter tungsten rod (A-M Systems) while in CMF PBS. The epithelium peels off readily as the tissue is filleted open, and the resulting slabs of mesenchyme were transferred by glass Pasteur pipette to Corning polycarbonate transwells in 6 well plates with the previously containing epithelial side of the mesenchyme facing up. To promote mesenchyme survival and morphogenesis, mesenchymal slabs were cultured in the presence of 1µM SAG. For tissue recombination, epithelial cell types were isolated from their respective cultures, centrifuged and subsequently resuspended in a small volume (<100µl) such that they were in a high-density slurry. To isolate epithelia from intestinal organoids (i.e., enteroids) or colonoids, organoids embedded within matrigel were rapidly pipetted to dislodge the organoid-matrigel composite. The organoid-matrigel mixture was subsequently centrifuged, resuspended in 1mL of basal media and the organoids were fragmented manually by pipetting the 1mL organoid suspended in basal media 20x through a 10µl pipette tip attached to a 1000µl tip. After centrifugation, the organoids fragments were resuspended to generate a cell slurry. To isolate human intestinal endoderm derived from cytoplasmic GFP-expressing pluripotent stem cell derived human intestinal organoids (HIOs), intact epithelial sheets were manually dissected from the loose mesenchyme of HIOs (that had been grown in matrigel for 30 days, see details below) before being transferred to the mouse mesenchyme. For isolation of MDCK cells, single cell slurry suspensions were generated by dissociating near-confluent cultures with TrypLE. For enteroids, colonoids, and MDCK recombinations, the cell slurries were then manually transferred by a P10 pipette to the isolated intestinal mesenchyme such that the mesenchymal explant tissues were overlayed completely by the dissociated epithelial cells, which in all cases rapidly formed cohesive sheets attached to the mesenchyme after 1 day of culturing. For isolated sheets of HIO-derived endoderm, the sheets were transferred directly by a glass Pasteur pipette onto the mouse mesenchymal slabs, and fine forceps were used to carefully position the sheets such that they lined the surface of the mesenchyme. We found that the human endoderm also remained as a cohesive sheet and quickly attached to the mesenchyme within 24 hours. Recombinant tissues were typically isolated following 72 hours of culture. Epithelial cell types used for recombination experiments were isolated, generated, and grown using standard protocols for mouse intestinal organoids and colonoids^99^, human pluripotent stem cell derived human intestinal organoids (HIOs)^23^ (and as outlined below), and MDCK cells^25^.

### Human pluripotent stem cell differentiation into intestinal organoids

Human embryonic stem cells (hESCs), H9 (WiCell Research Institute) and pluripotent stem cells, Wtc11-mEGFP (Coriell Institute/Allen Institute) were differentiated into human intestinal organoids as previously described^23,100^ with few modifications. Briefly, hPSCs (human pluripotent stem cells) were cultured with mTeSR^TM^ Plus medium (STEMCELL Technologies) on Geltrex^TM^ (Gibco) coated plates. To initiate definitive endoderm differentiation, hPSCs were first passaged using ReLeSR (STEMCELL Technologies) and plated onto Nunclon delta surface (Thermoscientific) 24-well plates priorly coated with Geltrex^TM^ (Gibco). Once cells reached 80% confluency, mTeSR Plus media was changed to definitive endoderm media (RPMI 1640, 1X GlutaMAX (Gibco), 100ng/ml Activin A and 2µM CHIR99021 (R&D Systems) on the first day of differentiation. Medium was changed for on day2-3 with definitive endoderm media without CHIR99021 but with 0.2-2% defined FBS (HyClone^TM^). To further differentiate the cells toward the hindgut, media was changed daily with hindgut medium (RPMI 1640, 1X GlutaMAX, 2% defined FBS, 2µM CHIR99021 and 500ng/ml FGF4 (R&D Systems) until day 8. On day 7 and 8, budding off spheroids were collected and replated in Geltrex onto 24-well plates with ENR intestinal growth media (Advanced DMEM/F-12, 1X B27 supplement (Gibco), 15mM HEPES (Gibco), 1X GlutaMAX, 1X Penicillin-Streptomycin (Gibco), 100ng/ml EGF (R&D Systems), 100ng/ml Noggin (R&D Systems) and 500ng/ml R-Spondin1 (R&D Systems)) to induce HIO formation. HIOs were split every 7-10 days.

### Transplantation of human intestinal organoids (HIOs)

HIOs were grown in ENR intestinal growth media for 27-34 days before transplantation into the kidney capsule of immunocompromised NOD-SCID IL2Rg null (NSG) mice (Jackson Laboratory strain no. 0005557) as previously described^101, 102^. 8-12 week male mice were used. Prior to transplantation HIOs were either: removed from Geltrex, washed with cold 1X PBS, incubated in Cell Recovery Solution (Corning) for 20 min on ice and resuspended in cold 1X PBS on the day of surgery. Alternatively, HIOs were embedded into purified Collagen 1 (Bio-Techne) and placed into regular ENR intestinal growth media overnight before surgery. Briefly, mice were anesthetized with 2% isoflurane, then an incision on the left flank was performed after shaving and sterilization with isopropyl alcohol and povidone-iodine to expose the kidney. One HIO was then placed into a subcapsular-created pocket and the kidney was returned to the peritoneal cavity. After closing the incisions, bupivacaine was administered at the site of surgery as a local anesthetic. Mice were euthanized and tHIO harvested 8 weeks after transplantation for immunofluorescence on cryosection or whole mount.

### Whole mount and section immunofluorescence

Tissues for whole mount or frozen sectioning immunofluorescence staining were fixed overnight at 4C in 4% PFA and subsequently washed twice for 10 minutes each in PBS. For whole mount staining, following PBS washes, tissues were cleared using a modified version of the CUBIC method^103^. First, tissues were incubated in 1.5mL Eppendorf tubes overnight in CUBIC-L solution on a rotator at 37C. Tissues were then washed twice for 5 minutes each in PBS. Tissue was then incubated in block/permeabilization solution (1X Animal Free Block Solution, 0.3% Triton X-100 in PBS) for 2 hours at room temperature or overnight at 4C. Tissues were next incubated in primary antibodies at specified dilutions in table below for at least two days at 4C in block/permeabilization solution. Following two, five minute washes in PBS, tissues were incubated in secondary antibodies with DAPI overnight at 4C. After another three washes in PBS to remove the secondary antibodies, tissues were incubated in CUBIC-R for at least one hour at room temperature and subsequently mounted on a slide using an image spacer (Grace Biolabs) so as not to crush the sample. For cryosectioning, tissues were fixed and washed as bove, transferred to 30% sucrose in PBS for 2 hours at room temperature or overnight at 4C, and subsequently incubated for 1 hour in OCT before being embedded in an OCT mold and sectioned at 12µm. tissue sections were stained using the above protocol without CUBIC treatment, and with primary antibody incubations of only one day. For EdU labeling, following immunofluorescent staining, EdU+ cells were detected using Click-iT Plus EdU Assay Kit (ThermoFisher, C10640). Images were taken on a Zeiss LSM900 confocal microscope. With clearing, the entirety of the tissue can be imaged. Where noted, optical slices, or side views refer to single plane images captured at the midplane of the gut from whole mount tissues, whereas top-down views refer to maximum intensity projections of the subepithelial layers imaged *en face* in whole mount tissues.

Immunofluorescence Antibodies.

**Immunofluorescence Antibodies**
| **Antibody (dilution)** | **Source** | **Catalog Number** |
| --- | --- | --- |
| CDH1 (1:300) | Cell Signaling Technology | #3195 |
| CDH1 (1:300) | BD Bioscience | #610182 |
| PDGFRA (1:500) | R&D Systems | #AF1062 |
| PDGFRA (1:300) | R&D Systems | #AF-307-NA |
| MYH9 (1:100) | BioLegend | #909901 |
| MYH10 (1:100) | BioLegend | #909802 |
| MYH14 (1:100) | Cell Signaling Technology | #8189s |
| pMLC (1:100) | abcam | #ab2480 |
| MYLK (1:50) | Sigma | #SAB4200808 |
| Fibronectin (1:300) | Abcam | #ab2413 |
| Collagen IV (1:100) | Bio-Rad | #2150-1470 |
| CDH4 (1:50) | R&D Systems | #AF6677SP |
| CDH11 (1:50) | ThermoFisher | #71-7600 |
| Phospho-Histone H3 (1:500) | Sigma | #06570 |

### Quantification of aggregate geometry and pattern

To measure aggregate aspect ratios, whole mount tissues were imaged *as* described above and the aspect ratio for each aggregate was calculated by first thresholding on the *PDGFRA^H2BGFP^* signal to set the boundaries of aggregates. Optical sections used for quantification were chosen at the midplane of each aggregate, equivalent to the maximum point of deformation in the overlying epithelium. After applying a Gaussian blur, the “Analyze Particles” feature in ImageJ/Fiji was used to identify aggregates (excluding sizes smaller than aggregates to avoid quantifying single or dispersed cells). Next, the region of interest defining each aggregate was fit to an ellipse, and the aspect ratio was calculated for each. To quantify aggregate pattern, whole mount stained tissues were fileted open, flattened with a coverslip, and imaged top-down (*en face*). After post-processing as above to identify aggregates, a nearest-neighbor distance calculation was performed. Equivalent segments of the tissue within the proximal jejunum (where cluster formation was robust after 48 hours of culture from E13.5) were used to compare between different condition.

### Single cell sequencing

*Pdgfra^H2BGFP^* mice on a C57BL/6J were used for the experiments. Small intestines were dissected from E13.5 – E18.5 embryos and cut into 2.5mm length segments along the proximal-distal axis. Number of tissues used and segments obtained from each tissue is summarized in the table below. Tissue segments were dissociated to single cells in CMF HBSS buffer containing 10mM HEPES, 1% BSA, 0.3mg/mL DNase and 0.1W/mL Liberase TL on an orbital shaker at 37C for 30 min with trituration by a P200 pipette every 15 mins. Single-cell suspensions were then transferred to 96-well U-bottom plates and washed twice in fetal gut media to remove any remaining BSA. MULTI-seq lipid barcode labeling was performed in 96 well plates following the MULTI-seq protocol^28^. Once the cells are incubated with lipid modified oligos (barcode, anchor and co-anchor) in total of 100µL volume reaction, 150µL of HBSS with 2% BSA solution were added to quench the reactions, which were subsequently pooled together in 15mL conical tubes following barcode labeling. The cells were pelleted and stained with APC/Cy7 anti-mouse Ter119, APC anti-mouse CD31, BV605 anti-mouse CD45, PE/Cy7 Epcam (all from BioLegend) for 30min on ice followed by a wash with FACS buffer (HBSS + 3% FBS + 10mM Hepes + 5mM EDTA + 1x P/S). The cells were resuspended in FACS buffer with DAPI to stain dead cells and passed through a 35µm cell strainer. Using a BD FACS Aria II, live (DAPI-, Ter119-) cells as well as enriched rare cells (CD31+, CD45+, Epcam+, or *Pdgfra^H2BGFP^*-high) were sorted separately. 35,000 cells were loaded to each well of GEM chip (2 wells of live-sorted cells and 1 well of enriched rare cells for each experimental day) for single-cell capture using the 10x Chromium Single Cell 3’ Reagent Kit V3 (10X Genomics) for aimed target recovery of 20,000 cells per well. Library preparation for both cDNA and MULTI-seq barcodes were performed following 10X library preparation protocol along with MULTI-seq library preparation protocol^28^. Libraries were pooled and subjected to sequencing in a single lane of an Illumina NovaSeq6000 for each experiment. Sequencing data was processed, and downstream analysis performed as described below.

Summary of tissue segments and replicates obtained from each tissue for the single-cell sequencing experiment.

| Stage | # Segments | # Replicates | Collection day |
| --- | --- | --- | --- |
| E13.5 | 3 | 4 | 2 |
| E14.5 | 5 | 3 | 2 |
| E15.5 | 10 | 2 | 1 |
| E16.5 | 15 | 2 | 1 |
| E18.5 | 20 | 2 | 2 |

### Processing FASTQ Files, assignment of barcodes, quality control and doublet Removal

Expression library FASTQs were pre-processed using cellranger count (10X Genomics) and aligned to the mm10 mouse reference transcriptome using default parameters. Read-depth normalization between 10x Genomics lanes was then performed using cellranger aggr (10x Genomics). Aggregated raw RNA UMI count matrices were then analyzed using Seurat^104^, excluding cells with < 1e3 total RNA UMIs and genes with ≤ 3 total counts across cells. Cells were further filtered to remove gene expression clusters associated with high proportions of mitochondrial reads and/or low total RNA UMIs. Using this filtered set of cell barcodes, MULTI-seq FASTQs were then pre-processed and subjected to the sample classification and semi-supervised negative cell reclassification workflows as implemented in the deMULTIplex R package (https://github.com/chris-mcginnis-ucsf/MULTI-seq). MULTI-seq sample classifications were then mapped onto gene expression space to identify clusters enriched for inter-sample doublets which were cross-validated using DoubletFinder^105^. The final scRNA-seq dataset was then generated by filtering cells using the final criteria: removing cells that were (i) classified by MULTI-seq as doublets or negatives, (ii) assigned to doublet-enriched clusters, or (iii) exhibited >= 10% mitochondrial gene content.

### Data integration, cell annotation and downstream analysis

Seurat objects were normalized using SCTransform methods^106^. Integration of Seurat objects from two different experimental days were performed following the Seurat integration workflow. Cells and clustering were visualized using Uniform Manifold Approximation and Projection (UMAP) dimensional reduction methods. To further analyze cells in specific clusters of interests, identified clusters were subsetted, integrated and re-clustered for downstream analysis and visualization. Cluster annotation was achieved by presence of known markers and based on identified marker genes. To identify marker genes of PDGFRa-High 1 population in the subepithelial mesenchymal cell population, differentially expressed genes were identified by comparing PDGFRa-High 1 to PDGFRa-low 1 population. The list of differentially expressed genes were then analyzed for functional enrichment analysis using EnrichR^107^ to identify upregulated KEGG and GO Terms. The Reactome database^108^ was used to compare PDGFRa-High progenitor population of the PDGFRa-low 1 populations further identify upregulated pathways at the bifurcation in cell types. Consensus Non-Negative Matrix Factorization (cNMF)^32^ and WOT analysis^33^ were applied as described (https://github.com/dylkot/cNMF and https://broadinstitute.github.io/wot/tutorial/) to further characterize the PDGFRA lineage. To generate the “Adhesion Score”, the list of PDGFRA^High^ driver genes defined by WOT analysis was reanalyzed using GO term analysis, which identified a subset of genes associated with the regulation of cell adhesion and cell-matrix adhesion (**Supplementary Table 6**) as being highly upregulated in this lineage. The collective expression of these gene module was plotted on the UMAP in Figure 2D. Scaled pathway activity score for pathways identified by the Reactome database was computed using the AUCell package^109^. Cell population density plots in Figure 2B were generated by overlaying subsets of data from 10x live-sorted lanes (blue) and FACS enriched rare cells lane, composed of PDGFRA^High^ cells (red) from respective developmental days on the overall population UMAP plot using stat_density2d function in ggplot. UMAP FeaturePlots were generated using scCustomize^110^.

### Laser ablation

Freshly harvested E14.5 tissue with *Pdgfra^H2BGFP^*, *R26R^mT/mG^* genotypes were used for the experiments for fluorescent visualization. Intestines were subject to the same epithelial removal protocol as described above. Following this, explanted mesenchyme was transferred to 35mm glass bottom dishes containing a thin coat of liquid 1% low melt agarose with the epithelial side of the mesenchyme facing up. The tissue was kept at the top of the agarose using forceps so that it remained mounted to the dish but was not embedded within the agarose and remained at the air-liquid interface. The tissue-containing plate was placed on a temperature controlled stage for the duration of the experiment and covered with fetal gut media containing 70µM nifedipine to block tissue movement due to smooth muscle contractility. A Nikon A1R upright laser scanning confocal equipped with a Mai Tai Deep See two-photon laser with Apo LWD 25x/1.1 water-immersion objective was used to laser ablate the top layer of mesenchyme that is normally adjacent to the now removed epithelium. 800nm wavelength with maximum (100%) laser power (1.8W with <100fs pulse width) with 1 frame per second scanning speed was used to make ablation on targeted regions of interest (ROI) by exposing this ROI over the period of 1 minute. Immediately after the 1 minute ablation period, the entire tissue region was imaged after switching to 15% laser power and the length of the cut was measured at this point and compared to the width of the ROI used to make the cut to generate the plots in Figure 4C.

### Microtissue spheroid experiments

To generate microtissue spheroids for rounding and coalescence experiments, intestinal mesenchyme from E14.5 embryos was isolated from the epithelium as described above and cultured in the presence of 1µM SAG for 3 days. During this time frame, distinct PDGFRA^High^ aggregates and PDGFRA^Low^ non-aggregates become clearly separated at the tissue surface. At this stage, these distinct regions were microdissected from the tissue using fine forceps and 0.004in diameter tungsten rods. The isolated tissue fragments were transferred to 96 well ultra-low attachment round-bottom plates (Corning) and cultured in DMEM with 10%FBS and 1µM SAG for 24 hours. For spheroid rounding experiments, the microtissues were imaged on a Zeiss spinning disk confocal to capture differences in rounding dynamics between the distinct populations. While PDGFRA^High^ aggregates round faster than PDGFRA^Low^ aggregates, they both generate spherical spheroids after 24 hours.

To perform spheroid coalescence experiments, the rounded microtissues were collected and placed pair-wise into a new 96 well ultra-low attachment round-bottom plate. To facilitate spheroid coalescence, the plate was subsequently centrifuged and a tungsten rod was used to manually put spheroids into contact. Spheroid coalescence was visualized on a Zeiss spinning disk confocal.

To perform PDGFRA^High^/PDGFRA^Low^ phase separation experiments, microtissues were isolated as above but from distal tissues that had yet to begin aggregating after 2 days of explant culture following epithelial removal, taking care to dissect the top layers of mesenchyme while leaving the muscle layers behind. The microtissue was allowed to round overnight in media containing 1µM SAG. Images were taken daily for consecutive days to capture separation of the two populations of cells.

### Measurement of ECM alignment

Following whole mount immunofluorescence in cleared tissues as described above, images of tissues stained for Fibronectin were acquired on a Zeiss LSM 900 and optical sections were acquired from specifically the subepithelial mesenchyme containing the PDGFRA^High^ cells. OrientationJ^111^ was then used to quantify Fibronectin/ECM alignment within a 100µm x 100µm region of a 5µm maximum intensity projection of the surface, subepithelial mesenchyme in either freshly isolated proximal or distal tissues or in tissues immediately following explant culture. ECM alignment parallel to the longitudinal (i.e., proximal-distal) axis of the gut was considered 0° while perpendicular to the longitudinal axis (i.e., circular alignment) was considered 90° as previously described using this same method^27^.

### Micropipette Aspiration

Micropipette aspiration was performed with pulled micropipettes generated from fire polished aluminosilicate glass capillary tubes (1.0mm outer diameter, 0.64mm inner diameter), and a ceramic tile was used to score the glass to generate 30µm-wide tips. Intestinal tissues from *Pdgfra^H2BGFP^*mice used for the experiment were harvested at E15.5, and following epithelial removal were cultured overnight in 1µM SAG to maintain aggregates. The explants were then cut into fragments and transferred to a rectangular well mounted on a coverslip such that the tissue fragments were suspended in fetal gut media containing 70µM nifedipine to prevent smooth muscle contractility. A micromanipulator was used to position the pipette and generate a seal on the tissue fragments. A constant pressure of ∼1kPa was applied to tissue fragments for 8 minutes. Image analysis was performed in Fiji, and deformability was measured as the length of maximum aspirated tissue at 8 minutes vs. the baseline position at the initial timepoint. PDGFRA^High^ vs. PDGFRA^Low^ tissues were confirmed by fluorescence intensity.

### Simulations

See **Supplementary Text** for detailed explanation and methods.

### Data and Code Availability

Original codes for the C-H and SPV models of mesenchymal aggregation are available at https://github.com/tjhakkin/GutCH and https://github.com/Gartner-Lab/GutSPV.

Raw scRNA-seq/MULTI-seq data are available through the following GEO Accession number: GSE233407 All other primary data or additional information required to reanalyze data reported here are available upon request.

## ACKNOWLEDGEMENTS

We are grateful for all members of the Gartner Lab, Klein lab, Jeremy Green, and the CZI Biohub Theory Group for their helpful feedback and suggestions on the project. We are additionally grateful for constant support from Pauline Marangoni, as well as Aimee Cortez, Evelyn Sandoval, Kelly Pan, Hadis Najibi, and for assistance with mouse maintenance and genotyping. We also thank the UCSF Biological Imaging Development Colab (BIDC), and the Parnassus Flow Core – part of the UCSF CoLabs – for providing essential equipment for these studies. This work was funded by R01DK126376, R35DE026602, F32DK128949, U01CA244109, R01GM135462, and by U01DK103147 from the Intestinal Stem Cell Consortium (ISCC), a collaborative research project funded by the NIDDK and the NIAID. This work was further supported by funding for the UCSF Center for Cellular Construction (CCC) by NSF. ZJG is an investigator of the Chan Zuckerberg Biohub San Francisco.

## CONTRIBUTIONS

TRH, HM, TJH, ZJG, and ODK conceived of the project. TRH wrote the manuscript with assistance from TJH, input from HM, ZJG, and ODK, and feedback from the other authors. TRH and HM performed the experiments and data analysis with assistance from EB, VS, and AK. TJH performed the computational modeling with assistance from JCS, MT, and KG and input from TRH and ZJG. DB, DV, CSM, and QZ assisted in the processing and analysis of the scRNA-seq data. Key resources were provided by HJ and WFD.

## SUPPLEMENTAL FIGURES

**Figure S1.**
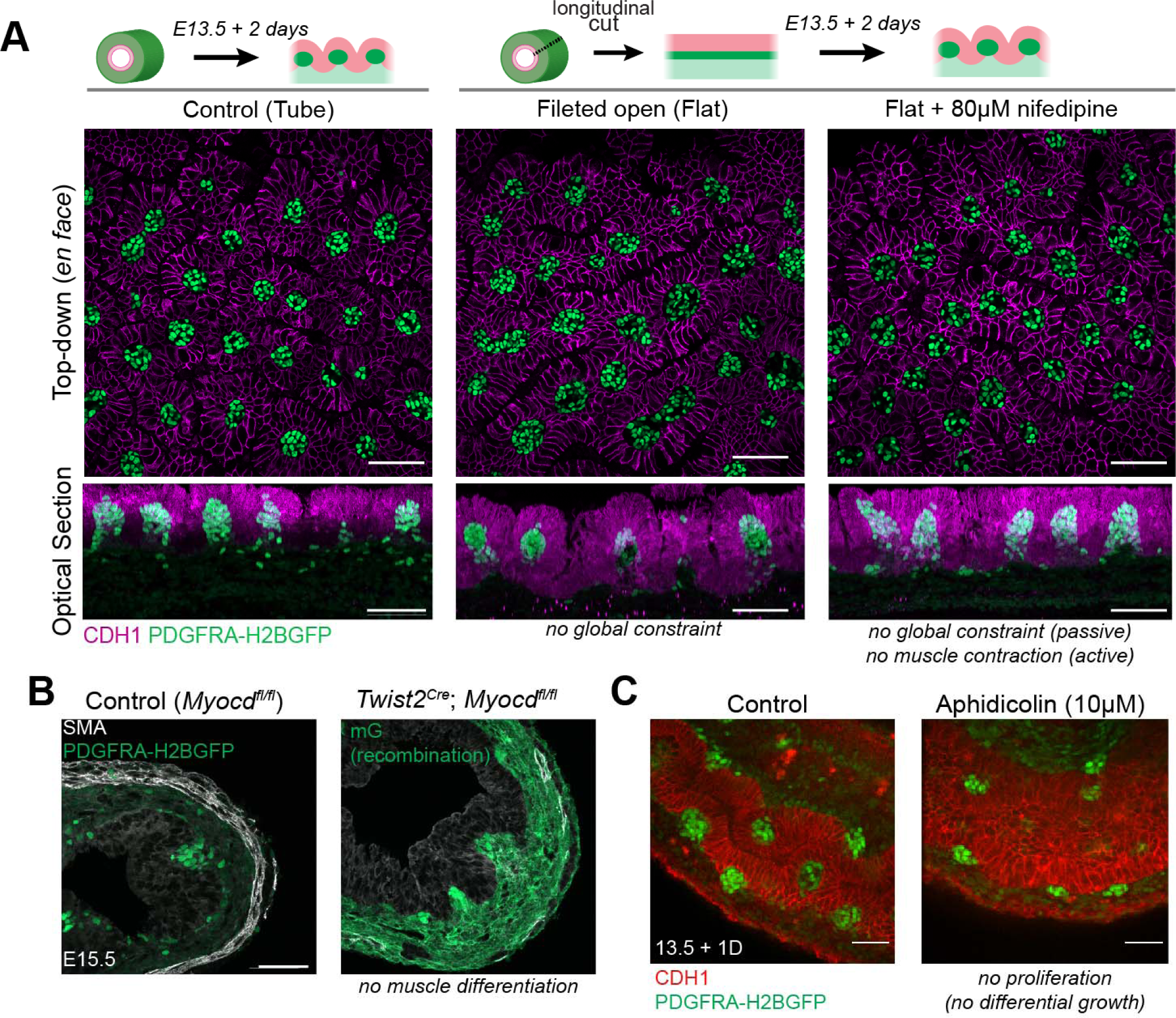
Smooth muscle constraint and proliferation are not required for mesenchymal aggregation, related to Figure 1. (A) Top-down (*en face*) views of a whole gut grown as a tube (control), fileted open (i.e., flat, non tubular), or fileted open and cultured in the presence of nifedipine to inhibit smooth muscle contractility for 2 days starting from E13.5 when no clusters are present. Bottom panels are orthogonal reslice/reconstructions from whole mount image in the top panels and highlight that under all conditions, mesenchymal cells aggregate and epithelial folding occurs. (B) Cross sections through E15.5 small intestine of control mice or those with *Myocd*, a master regulator of smooth muscle differentiation, conditionally knockout out specifically in the mesenchyme by *Twist2^Cre^*. Smooth muscle differentiation within the circumferential and longitudinal layers is absent in these mutants in the proximal intestine (imaged here), but mesenchymal aggregates still form and deform the overlying epithelium as in controls. Slight staining for alpha-smooth muscle actin (SMA) present in the mutants is visible in vascular presumptive vascular smooth muscle. Recombination by *Twist2^Cre^,* which is broadly expressed in the mesenchyme, is noted by green membrane via the recombination marker *R26R^mT/mG^*, while the *PDGFRA^H2BGFP^*reporter is nuclear. (C) Optical sections through whole mount tissue. Aphidicolin, a proliferation inhibitor, does not impair the mesenchymal aggregation or the initiation of tissue curvature. Scale bars = 50 µm

**Figure S2.**
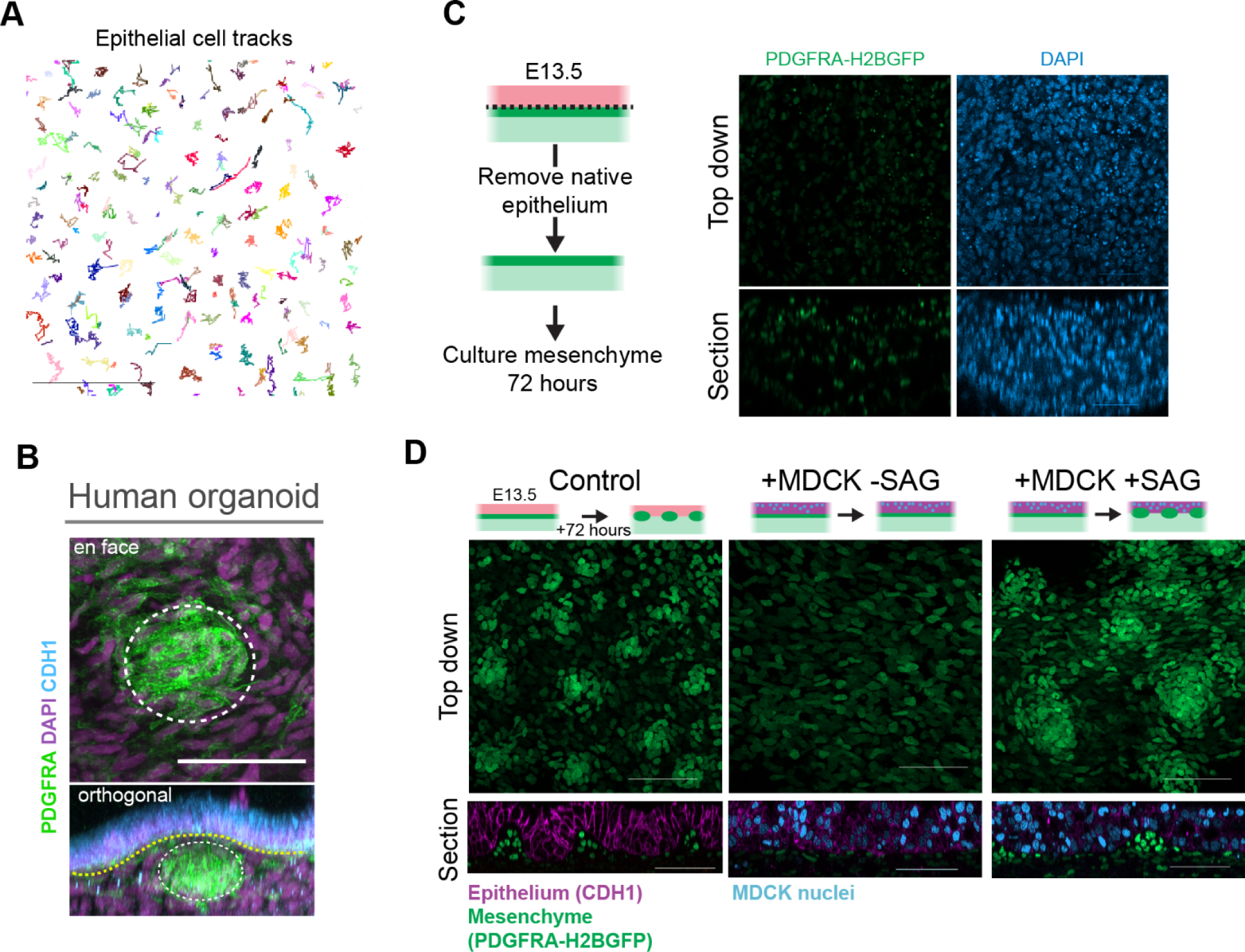
Hedgehog signaling induces mesenchymal dynamics sufficient for aggregation, related to Figure 1. (A) Representative epithelial cell tracks from a timelapse movie of the E14 small intestine over 16 hours. Note lack of neighbor exchange and largely constrained movement. Epithelial cells are labeled by the *R26R^nucMkate^*^2^ reporter. (B) Optical sections from a transplanted Human Intestinal Organoid isolated after 8 weeks and stained and imaged in whole mount to capture the 3D structure of the aggregate and tissue curvature. Aggregates appeared coincident with and immediately beneath sites of epithelial folding. (C) Whole mount top down view and orthogonally reconstructed sections from a mesenchyme-only tissue cultured in basal media from E13.5 for three days. Note lack of PDGFRA^High^ cells and puncta in DAPI channel indicative of cell death. (D) Representative images of small intestinal explant tissues cultured in various conditions. While the mesenchyme survives in the presence of the MDCK epithelial layer, mesenchymal dynamics are not induced unless Hedgehog signaling is activated by SAG. Scale bars = 50 µm

**Figure S3.**
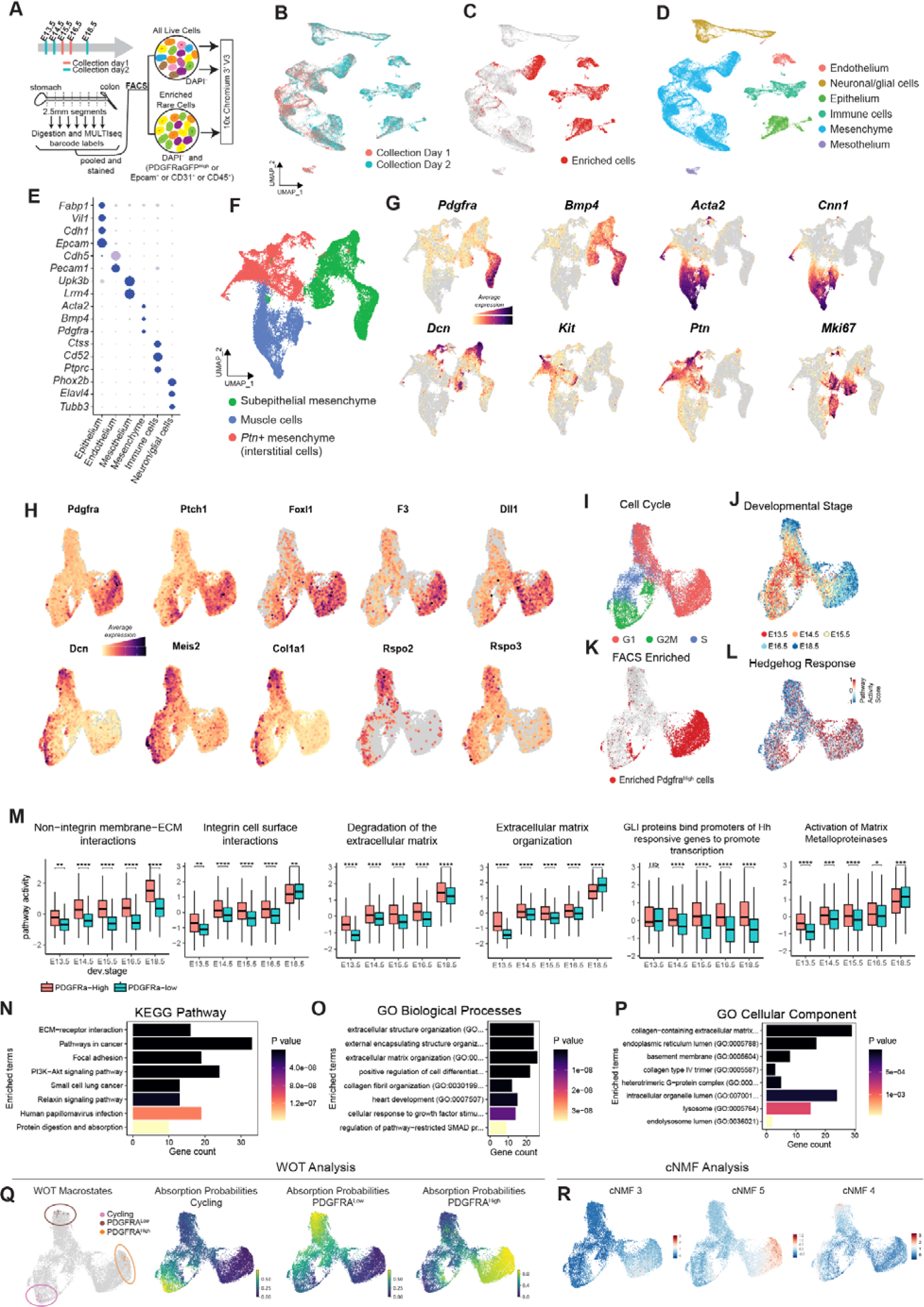
Differential gene expression during intestinal mesenchymal development, related to Figure 2. (A) Schematic summarizing experimental setup. E13.5 - E18.5 *PDGFRA^H2BGFP^* intestinal tissues were collected over two different experiments. Tissues were cut into 2.5 mm pieces, digested and labeled with MULTI-seq lipid-modified barcodes. Unmodified proportions of all live cells and populations enriched for PDGFRA^High^ cells, immune cells, endothelial cells and epithelial cells were sorted and captured for single-cell sequencing using 10x Chromium V3 kits. (B) Overlays of data collected in two different batches reveal successful integration of data. (C) Cell populations enriched by FACS map to the expected cluster in following panel. (D) UMAP projection of all cells collected, clustered, with major cell types identified by expression of known marker genes. (E) Dot plot of key marker genes used for cluster cell type identification (F) UMAP projection and reclustering of the mesenchymal cell subset, highlighting three major categories of mesenchymal cell types identified by marker gene expression in the following panel. (G) UMAP expression profiles of cluster-defining marker genes. (H) UMAP projections of the subsetted and reclustered subepithelial mesenchyme and presumptive cycling progenitors as outlined in Figure 2, based on expression of known subepithelial marker genes. Expression profiles show additional markers of the PDGFRA^High^ and PDGFRA^Low^ cells previously associated with intestinal development or function. (I) Cell cycle stages projected onto the UMAP. (J) Developmental stages of tissue collection projected onto the UMAP show prospective temporal trajectories for each lineage. (K) Overlay of PDGFRA^High^ cells in enriched by FACS shows good agreement with *Pdgfra* expression. (L) Reactome pathway score of “GLI proteins bind promoters of Hh responsive genes to promote transcription” plotted on the UMAP shows enrichment in the PDGFRA^High^ cells. (M) Box plots for Reactome pathway activity levels at each stage of development. See **Supplemental File 3** for complete pathway activity lists and *p* values. (N) Top 10 KEGG Pathways enriched in PDGFRA^High^ vs PDGFRA^Low^ cells. (O) Top 10 GO Biological Processes enriched in PDGFRA^High^ vs PDGFRA^Low^ cells. (P) Top 10 GO Cellular Components enriched in PDGFRA^High^ vs PDGFRA^Low^ cells. (Q) Three macrostates are calculated by WOT analysis and are mapped onto the UMAP. These macrostates correspond to the endpoint of temporal differentiation trajectories into PDGFRA^High^, PDGFRA^Low^ and cycling cell states. Accompanying absorption probability plots show the calculated probability of each cell within the subepithelial mesenchyme to align with each of the three macrostates, representing the prospective differentiation trajectories of each lineage. See **Supplemental File 4** for corresponding gene lists. (R) UMAP showing two gene expression programs (composed of 250 genes each) identified by consensus non-negative matrix factorization (cNMF) to contribute to the PDGFRA^High^ cell state (cNMF 3 and 5) and one contributing to the PDGFRA^Low^ state cNMF 4). See **Supplemental File 5** for corresponding gene lists.

**Figure S4.**
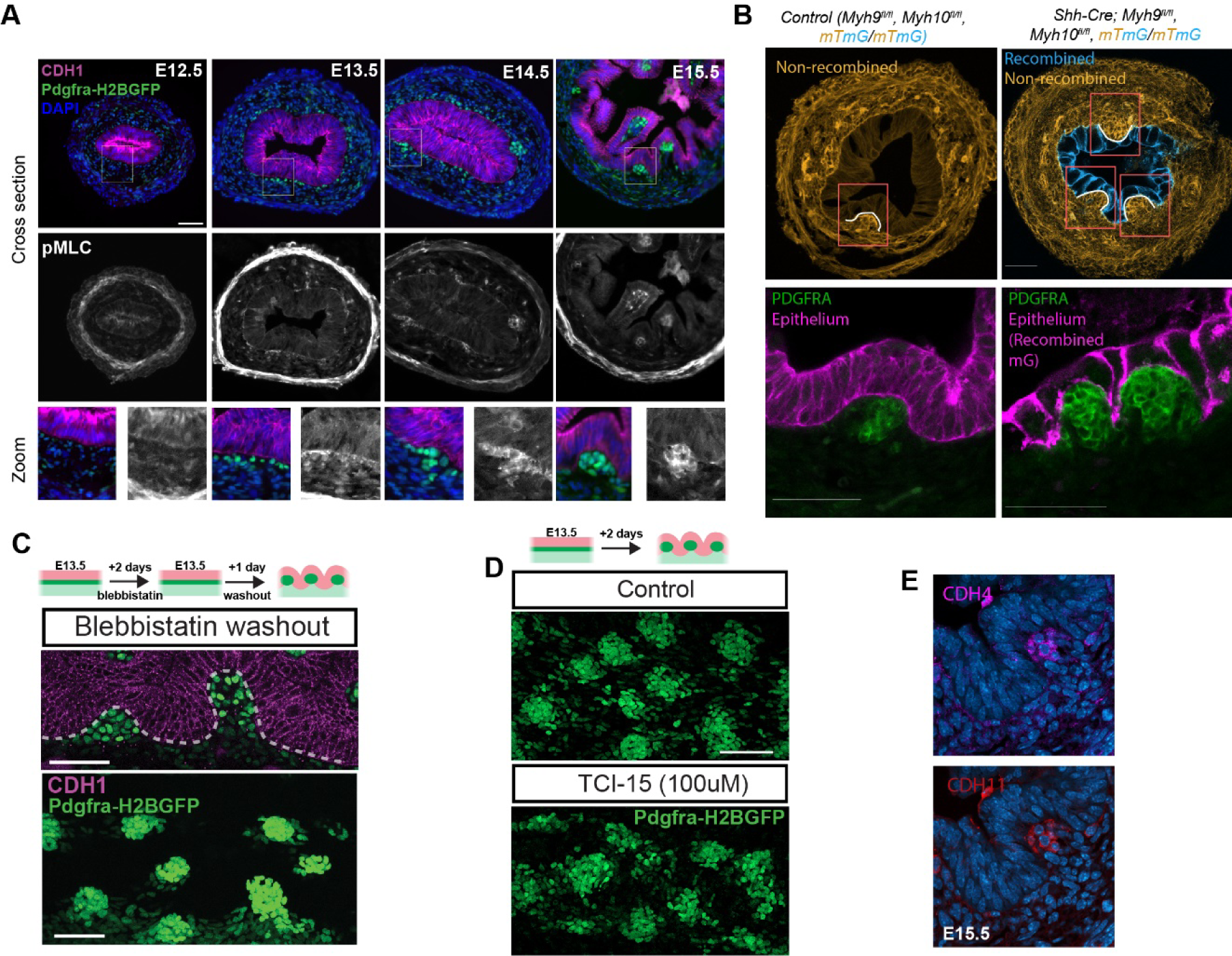
Mesenchymal actomyosin activity and cell adhesion underly aggregate formation, related to Figure 3. (A) Expression of phospho-myosin light chain (pMLC) over developmental time in small intestine tissue cross sections. pMLC emerges coincident with PDGFRA^High^ cells in the subepithelial mesenchyme. Note that it is also highly expressed in the circumferential smooth muscle layer as expected. pMLC staining is relatively lower or absent in the epithelium. Bottom panels show higher magnifications of the tissue interface. (B) Tissue cross sections through distal E15.5 intestines in control and mutant guts lacking *Myh9* and *Myh10* expression in the epithelium. While epithelial cell morphology is grossly abnormal, with relatively large and few cells, mesenchymal clustering is not impaired, and epithelial deformation along the basal surface of the tissue interface is still apparent immediately above aggregates. Boxed regions highlight sites of aggregates and folding in the mutant. Bottom panels show magnified expression of PDGFRA stained in separate samples from those in the Top. (C) Tissue cross section and whole mount from an intestinal explant treated with blebbistatin for 2 days, after which it was washed out and replaced with normal media, highlighting the reversibility of blebbistatin inhibition on tissue morphogenesis. (D) Whole mount top down views of clusters in control tissue explants and those treated with the A2B1 integrin inhibitor TCI-15, which alters and generates a higher variability in cluster dimensions (see Figures 3 and 5 for quantification). (E) IF is tissue cross sections of the small intestine at E15.5 highlighting the localization of CDH4 and CDH11 to cell interfaces within presumptive mesenchymal aggregates at the tips of emerging villi, but is not visible in regions that are not aggregating. Scale bars = 50 µm

**Figure S5.**
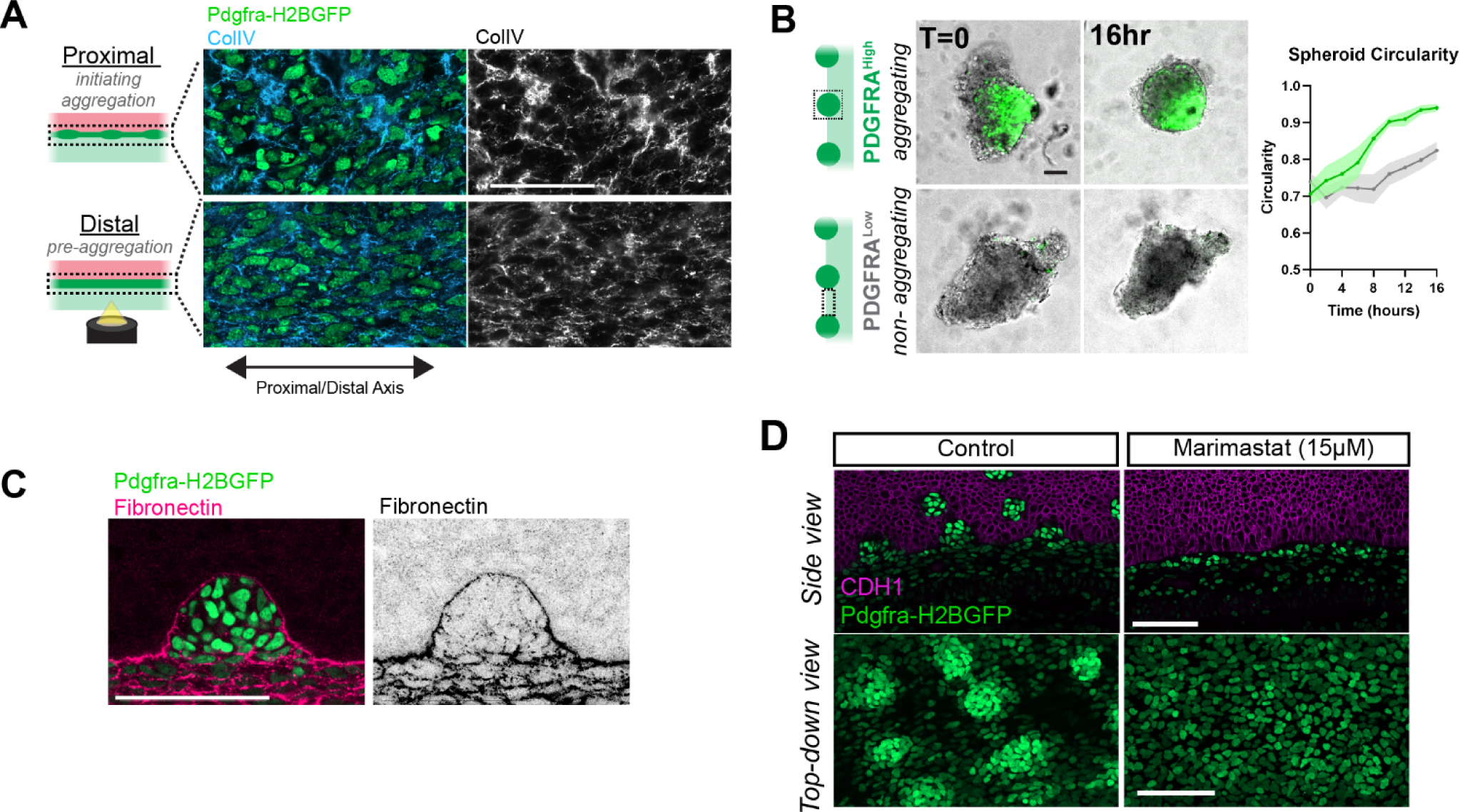
Matrix remodeling promotes tissue fluidity, related to Figure 4. (A) Whole mount optical sections focused at the subepithelial mesenchyme with proximal/distal axis aligned horizontally. IF for Type IV Collagen. (B) Microtissue spheroid rounding assay. PDGFRA^High^ tissues round faster than PDGFRA^Low^ tissues. are mean ± SD from *n =* 8 PDGFRA^High^ spheroids and *n =* 12 PDGFRA^Low^ spheroids. (C) Optical cross section at E15.5 demonstrating reduced fibronectin expression within the aggregate, consistent with matrix degradation and remodeling in the PDGFRA^High^ tissue phase. (D) Images from explanted intestinal tissues treated for 2 days from E13.5 with the broad spectrum MMP inhibitor marimastat demonstrating a failure of mesenchymal aggregation but persistence of PDGFRA^High^ differentiation at the epithelial-mesenchymal interface. Scale bars = 50 µm

**Figure S6.**
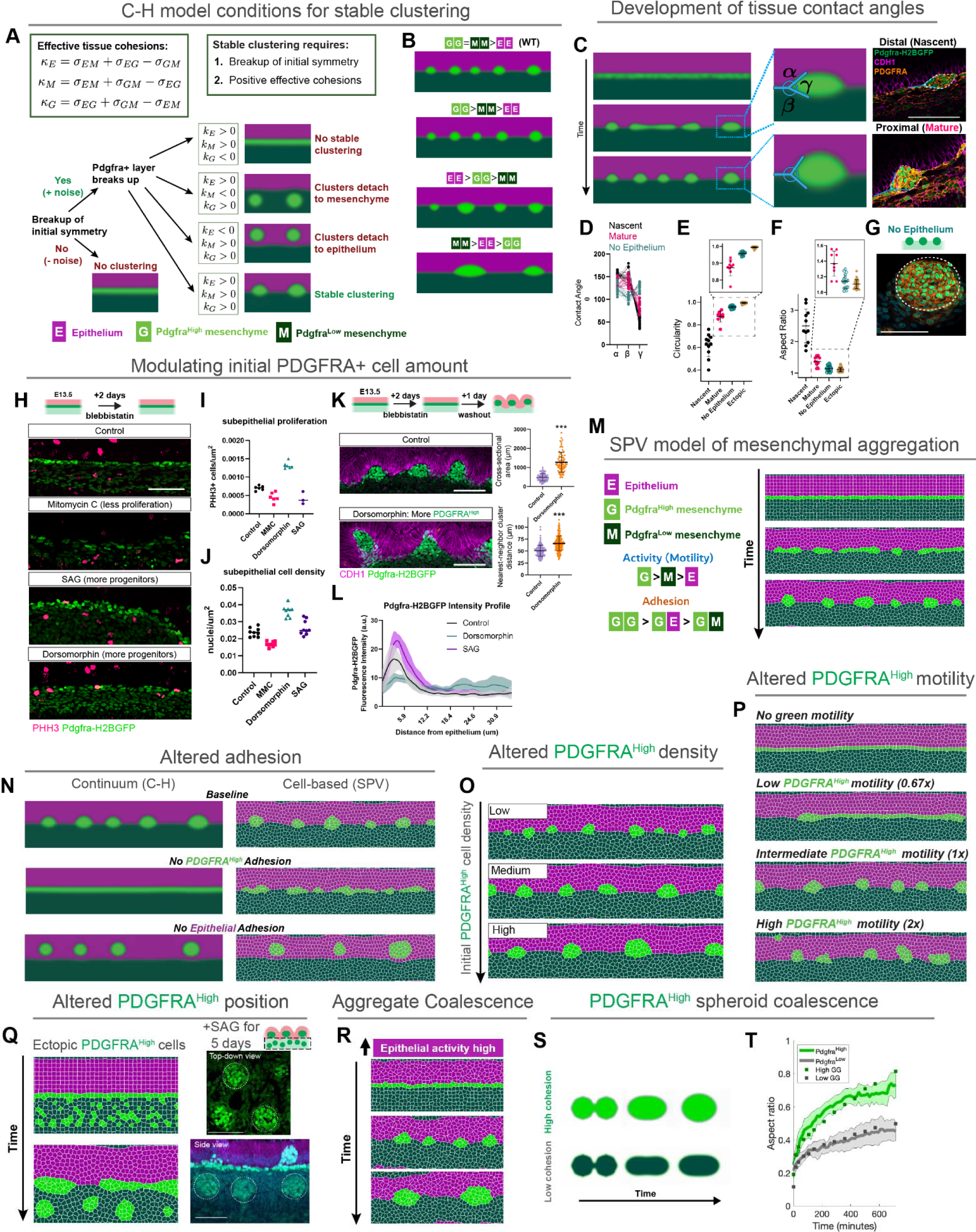
Computation models of cell aggregation and interfacial folding, related to Figure 5. (A) Flow chart defining the outcomes of the C-H model when run with different parameters. Without noise, no clustering is observed. With noise in the PDGFRA^High^ layer, aggregates form unless the effective cohesion of the PDGFRA^High^ phase is negative. The position of aggregate formation is dependent upon the effective cohesion of the other layers. Only when the effective cohesion is positive for all three layers do aggregates form at the interface. (B) Effective of varying cohesion hierarchies on PDGFRA^High^ aggregate morphology. Model is run with all positive effective cohesion values. Droplet morphology and contact angles with the surrounding interfaces is dependent upon this hierarchy. (C) Evolution of tissue contact angles over developmental time when model is run as WT. When dewetting from the surrounding interfaces, the PDGFRA^High^ aggregates develop characteristic contact angles that mature over time. Images are representative from WT tissues highlighting distinctive contact angles of the aggregates in mature (more proximal) versus nascent (more distal) clusters in an E14.5 intestine. (D) Measurement of *in vivo* contact angles as in panel C. Contact angles are calculated for individual aggregates from three E14.5 intestines where clustering was visible in the proximal gut tissue. Mature and nascent clusters were defined by their position either at the proximal (mature) end of the wavefront of aggregation, or at the distal (nascent) end of the wavefront. Contact angle measurement was also performed on tissues that had their epithelia removed at E14.5 and cultured for 3 days. For epithelial free tissues, only aggregates that formed on the lateral sides of the tissue were measured, as those that formed at the midline were effectively compressed by the surface tension at the air-liquid interface of the culture setup. (E) Circularity measurements of the same aggregates as in panel D, as well as ectopic deep aggregates generated by SAG treatment as in Figure 5F and epithelial free aggregates as in panel G. Measured at the midplane of the cluster. Magnified insets are shown above. (F) Aspect ratio of the same aggregates as in D, as well as ectopic deep aggregates generated by SAG treatment as in Figure 5F and epithelial free aggregates as in panel G. Measured at the midplane of the cluster. Magnified insets are shown above. (G) Example of spherical aggregate that forms with distinct geometries when the epithelial interface is removed. (H) Optical sections through explanted intestinal tissues at the epithelial mesenchymal interface immunostained for phospho-histone H3 to mark proliferation. Tissues were treated with labeled pharmacological agents for 2 days in the presence of blebbistatin, which pauses morphogenesis but allows for PDGFRA+ differentiation at the subepithelial interface. (I) Quantification of subepithelial PDGFRA+ cell proliferation from images as in panel H. Each point represents an image quantified from an independent explant. Dorsomorphin increases the number of PDGFRA+ proliferating cells at the tissue interface, while MMC decreases the number. (J) Quantification of subepithelial PDGFRA+ cell density, measured as the number of nuclei in a region of interest including the first 2 layers of cells adjacent to the epithelium in panel H. Dorsomorphin leads to an increase in PDGFRA cell density at the interface. (K) Optical section through control explant or explant treated with Dorsomorphin for 2 days in the presence of Blebbistatin followed by a 1 day washout. Cluster size and spacing are increased, quantified in plots on the right. Experiment was carried out simultaneously with samples in Figures 5C-E and as such use the same control data, re-graphed here for direct comparison. See Methods for additional details. (L) *PDGFRA^H2BGFP^* intensity line plots measured as distance away from the epithelium. SAG treatment expands the domain of PDGFRA^High^ cells within the subepithelial mesenchyme. (M) SPV (cell-based) simulation of PDGFRA^High^ layer breakup into aggregates. Outline of the three tissue layer SPV model recapitulating intestinal morphogenesis. Hierarchical parameters are outlined for the simulation shown, considered as control, wild-type simulations in following experiments. See **Supplemental Text** for further details. (N) SPV simulation with varying initial PDGFRA^High^ values demonstrating scaling of droplet size and spacing. (O) Side-by-side comparison of SPV and C-H simulations demonstrating similar outcomes over time. Unlike in the cell-based SPV model, where the noise is explicitly coded into the motility of the cell particles, in C-H the motility is implicitly present in the continuum material phases to reorganize through diffusion. (P) SPV simulations run with varying amounts of PDGFRA^High^ motility. (Q) SPV model run with PDGFRA^High^ cells patterned throughout the mesenchyme results in ectopic deep clusters, similar to tissues shown on right treated with SAG for 5 days. (R) SPV model run with high epithelial motility. In this condition the epithelium is essentially fluidized, and actively undergoes neighbor exchange. This lowers the barrier for the ability of aggregates at the interface to move, resulting in coalescence at shorter timescale than what would be observed in the wild-type simulation wherein the epithelium is static (more solid-like). (S) C-H simulation of microtissue spheroid coalescence of either PDGFRA^High^ spheroids (high cohesion) or PDGFRA^Low^ spheroids (lowered cohesion) as in Figure 4E. In this case the spheroids are represented as the individual phases and are initialized at the point when they come in contact. (T) Concordance between the simulation and experiment of spheroid coalescence from panel S and 4E. The lines represent the data (Same dataset as portrayed in Figure 4E) and the dots represent the model predictions. Scale bars = 50 µm

## SUPPLEMENTAL FILE INFORMATION

### Supplemental Videos

**Video S1. Co-emergence of epithelial folding and mesenchymal aggregation**

Timelapse video of a small intestinal explant from an E14 *Pdgfra^H2BGFP^; Shh^Cre^; R26R^tdTomato^*mouse. *Pdgfra^H2BGFP^-*labeled cells are in green, and epithelial cells, labeled by *Shh^Cre^; R26R^tdTomato^* are in magenta. The tissue has been fileted open and is viewed from the side to capture morphogenesis at the tissue interface. No aggregates were present at the initiation of the movie, but begin forming shortly thereafter. The emergence of aggregates and folding occurs simultaneously.

**Video S2. Mesenchymal aggregation of PDGFRA^High^ cells**

Timelapse video of a small intestinal explant from an E14 *Pdgfra^H2BGFP^*mouse capturing the onset of cell aggregation in the mesenchymal layer. Pdgfra^H2BGFP^signal is shown in inverted greyscale. Time is in hh:mm.

**Video S3. Diffusive-like cell motility during mesenchymal aggregation**

Timelapse video of a small intestinal explant from an E14 *Pdgfra^H2BGFP^; Twist2^Cre^; R26R^Confetti^*mouse, specifically focusing on the proximal region of the tissue at the onset of aggregation. Left panel shows merger of the *Pdgfra^H2BGFP^*signal in green and sparsely labeled mesenchymal cells (both PDGFRA^High^ and PDGFRA^Low^) are labeled with cytoplasmic RFP by*Twist2^Cre^; R26R^Confetti^* and shown in orange. The right panel shows the single RFP channel. The focal plane is specifically fixed on the first two layers of cells in the subepithelial mesenchyme containing the PDGFRA+ cells. Time is in hh:mm.

**Video S4. Restricted motility in the pre-aggregate tissue**

Same as S3, but the timelapse movie captures the distal small intestine where aggregation has yet to begin and does not occur during the timeframe of the video. Time is in hh:mm.

**Video S5. Continued cell motility and neighbor exchange with aggregates**

Timelapse video of a small intestinal explant from an E15 *Pdgfra^H2BGFP^; Twist2^Cre^; R26R^Confetti^* mouse, of aggregates that have already formed. Time is in hh:mm.

**Video S6. Aggregates are destabilized by integrin inhibition**

Timelapse videos intestinal explants from E14.5 *Pdgfra^H2BGFP^*mice (shown in inverted greyscale) incubated in the presence of 100µM TCI-15 to inhibit A2B1 integrin (right video) or control (left video). Aggregate position is highly stable in controls. Treatment with TCI-15 causes the aggregates to become destabilized, with clusters continuously forming and dissolving and cells moving between aggregates. Time is in hh:mm.

**Video 7. Aggregates coalesce in the absence of a restrictive epithelium**

Timelapse videos from small intestinal explants from E14.5 *Pdgfra^H2BGFP^*mice that had been cultured already for 2 days, following epithelial removal. Aggregates come into contact, fuse, and coalesce to form larger, but fewer aggregates. Time is in hh:mm.

### Supplemental Tables

**Supplemental Table 1. Transcriptional markers of subepithelial cell clusters**

Marker genes for each cell cluster within the subepithelial mesenchyme shown in Figure 2C, generated by the “FindAllMarkers” command in Seurat.

**Supplemental Table 2. Transcriptional markers of PDGFRA^High^ vs. PDGFRA^Low^ cells**

Marker genes differentially expressed by PDGFRA^High^ vs. PDGFRA^Low^ cells. Table generated by the “FindMarkers” command in Seurat, comparing genes expressed by the (combined) PDGFRa-High 1 cluster and PDGFRA-High progenitor versus the PDGFRa-low 1 cluster, represented as log fold change of PDGFRA^High^ vs. PDGFRA^Low^.

**Supplemental Table 3. Reactome pathway activity**

List of Reactome pathways and activities within the PDGFRA^High^ vs. PDGFRA^Low^ vs. Cycling Cell clusters.

**Supplemental Table 4. WOT analysis driver genes**

List of genes identified by WOT analysis to be drivers of the PDGFRA^High^, PDGFRA^Low^, or cycling macrostates.

**Supplemental Table 5. cNMF gene expression programs**

List of the top 250 genes contributing to the cNMF identities in Figure S3R.

**Supplemental Table 6. Adhesion Score**

List of the genes contributing to the Adhesion Score in Figure 2D

### Supplementary Text

**Supplementary Text. Description of the C-H and SPV models**

