## Supplementary Text for "Patterning and folding of intestinal villi by active mesenchymal dewetting"

**Cahn-Hilliard model for villus clustering**

We assume that during the critical period of tissue reorganization leading to the initial PDGFRA^High^ cluster formation the tissue layers can be approximated as behaving liquid-like, with viscous reorganization driven by differential tissue adhesions dominating over tissue elasticity. While elasticity has been proposed to play dominant role during the later stages of morphogenesis involving villus elongation^1^, our focus here is strictly at the primary cluster formation dynamics, prior to the onset of significant epithelial buckling or growth.

In the Cahn-Hilliard (C-H) framework we describe each tissue type $i$ as a material phase with concentration given at position $x$ (now in 2D) and time $t$ by function $c_{i}(x,t)$. Tissue reorganization is assumed to be driven by the global minimization of total tissue energy. To find an expression for the tissue energy, we first consider the adhesion energy density written as a simple triple-well potential

$\begin{aligned} g(\text{c})=g(c_{1},c_{2},c_{3})=\sigma_{12}c_{1}^{2}c_{2}^{2}+\sigma_{13}c_{1}^{2}c_{3}^{2}+\sigma_{23}c_{2}^{2}c_{3}^{2}, \end{aligned}$	(1)

where $\sigma_{ij}$ are the surface tensions (or energies) at tissue boundaries. While the specific indexing is not crucial, in the following it is assumed that the phases 1, 2, 3 correspond to epithelium, PDGFRA^Low^ and PDGFRA^High^, respectively. As a technical side-note, for practical simulations we augment Eq. (1) with higher order terms in order to improve the accuracy of the solution, which we will outline later. For the present purposes of introducing the key model parameters and dynamics, without loss of generality, the basic form of Eq. (1) suffices.

Because energy (1) cannot distinguish between different distributions of phase clusters over a tissue volume (e.g., many small clusters vs. few large), a gradient penalty term is added, giving the total energy density

$\begin{aligned} f(\text{c})=g(\text{c})+\epsilon^{2}\overset{3}{\underset{i=1}{\sum}}\frac{\kappa_{i}}{2}|\nabla c_{i}|^{2}, \end{aligned}$	(2)

where $\epsilon>0$ is a parameter proportional to the thickness of phase interfaces, and $\kappa_{i}$ are obtained from the surface tensions $\sigma_{i}$ such that

$$\begin{matrix} \kappa_{1} & =\sigma_{12}+\sigma_{13}-\sigma_{23}, \\ \kappa_{2} & =\sigma_{12}+\sigma_{23}-\sigma_{13}, \\ \kappa_{3} & =\sigma_{13}+\sigma_{23}-\sigma_{12}. \end{matrix}$$

The values $\kappa_{i}$ can be obtained by equating a two-phase reduction of energy (2), i.e., $f(c,1-c,0)$, with a two-phase density $f_{2}(c)=\sigma c^{2}(1-c)^{2}+\epsilon^{2}\sigma|\nabla c|^{2}$, and solving for the values of $\kappa_{i}$. In a later section we will identify values $\kappa_{i}$ as the effective tissue cohesions, which are the key parameters controlling the clustering dynamics in the continuum model.

The total tissue energy is obtained by integrating the energy density (2) over the whole tissue $\Omega$

$$\begin{aligned} F(\text{c})=\int_{\Omega}fdV. \end{aligned}$$

Using standard variational techniques we find that the global minimizer condition for $F(\text{c})$ is

$\begin{matrix} \mu_{i}=\frac{\delta F(\text{c})}{\delta c_{i}}=\frac{\partial g(\text{c})}{\partial c_{i}}+\epsilon^{2}k_{i}\nabla^{2}c_{i}=0 & & in\Omega, \end{matrix}$	(3)

where $\mu_{i}$ is commonly called the chemical potential of phase $i$.

According to the Fick's first law, the flux of particles in a system can be expressed as proportional to the chemical potential

$$\begin{aligned} \text{J}_{i}=-M_{i}\nabla\mu_{i}, \end{aligned}$$

where $M_{i}$ is the mass mobility of phase $i$. Combining this with the conservation of mass, we obtain the Fick's second law for material diffusion

$$\begin{aligned} \frac{\partial c_{i}}{\partial t}=-\nabla\cdot\text{J}_{i}=\nabla\cdot(M_{i}\nabla\mu_{i}), \end{aligned}$$

where $\mu_{i}$ are defined by Eq. (3). Two boundary conditions are usually ascribed: $\kappa\nabla c\cdot\text{n}=0$ on $\partial\Omega$ and $\nabla\mu\cdot n=0$ on $\partial\Omega$. In our implementation we set periodic boundaries for concentrations to avoid boundary effects on the clustering.

The C-H model was implemented in Python 3.10 using the DOLFIN 2019.2.0 library of the FEniCS project^4,5^ for the numerical solver. For implementation details and the parameter values used in the simulations of Figs. 5 and S6, see *https://github.com/tjhakkin/GutCH.*

**Consistent form of adhesion energy density**

Boyer et al.^2^ noted that using a simple triple-well potential of form Eq. (1), which is commonly encountered in literature, results in a model that is inconsistent with the corresponding two-phase case. The practical side-effects of using an inconsistent energy density, while numerically possible, include non-physical “leaking” of one phase into another along the phase boundaries, and formation of contact angles that are inconsistent with the values predicted directly from the surface tensions $\sigma_{i}$. As a remedy to these issues, Boyer et al. propose augmenting the energy density with higher order terms, specifically replacing the density of Eq. (1) with

$\begin{aligned} g(\text{c})=\sigma_{12}c_{1}^{2}c_{2}^{2}+\sigma_{13}c_{1}^{2}c_{3}^{2}+\sigma_{23}c_{2}^{2}c_{3}^{2}+c_{1}c_{2}c_{3}(\kappa_{1}c_{1}+\kappa_{2}c_{2}+\kappa_{3}c_{3})+3\Lambda c_{1}^{2}c_{2}^{2}c_{3}^{2}, \end{aligned}$	(4)

where $\Lambda$ is a positive constant whose exact value is not critical, provided it is large enough. In all our simulations we use $\Lambda=7$. Note that in the special case two two phases, Eq. (4) reduces into the standard double-well potential $g(c,1-c,0)=\sigma c^{2}(1-c^{2})$ commonly used in two-phase models.

**Effective tissue cohesion**

In the above we gave expression for the model parameters $\kappa_{i}$ as a function of surface tensions $\sigma_{i}$. In the literature scalar $S_{i}=-\kappa_{i}$ is commonly called the *spreading coefficient* of phase *i^2,3^*. We interpret $\kappa_{i}$ as representing the *effective cohesion* of phase $i$ by the following reasoning. Consider, for example, $\kappa_{1}=\sigma_{12}+\sigma_{13}-\sigma_{23}$: If, other things being equal, the costs of interfaces that phase 1 can form with phases 2 and 3 are high (large $\sigma_{12}$ and $\sigma_{13}$, respectively) then a high ratio of interior volume to surface area of phase 1 is favored, giving the phase a high effective cohesion. On the other hand, relatively high surface tension $\sigma_{23}$ suggest that phases 2 and 3 will avoid forming a common interface, driving them to promote the spreading of phase 1 along the interface, hence lowering the effective cohesion of phase 1.

For successful clustering in our case, that is, for stable clusters to emerge at the interface between the mesenchyme and the epithelium such that the clusters do not end up leaving the interfacial region, a necessary condition is that $\kappa_{i}>0$ for all $i$. In other words, all phases must have positive effective cohesion. Fig. S6A demonstrates what happens when one of $\kappa_{i}$ fail to satisfy this condition: If the $\kappa_{i}$ corresponding to the clustering phase is zero or negative, the clusters will fail to form by not having sufficient effective cohesion to drive rounding to clusters. If $\kappa_{i}$ of a neighboring phase is zero or negative, the clusters will be engulfed by that phase, detaching from the interfacial region; and while technically the clustering phase still forms clusters, we consider this failed clustering in practice, given that the scenario clearly does not correspond to proper observed clustering in tissues.

**Self-propelled Voronoi model**

Our self-propelled Voronoi (SPV) model implementation follows the methods present by Bi et al.^6^ and Barton et al.^7^, and we refer the reader to those studies for detailed discussion about the model. Briefly, in SPV the tissue domain is divided into polygons representing cells, with each polygon $i$ having a target area $A_{i}^{0}$ and target perimeter $P_{i}^{0}$. Each polygon edge $(\mu,v)$ of length $l_{\mu v}$, representing the boundary between two cells, is associated with a cost $\Lambda_{\mu v}$, which corresponds to the surface energy parameter $\sigma_{ij}$ for heterotypic (non-kind) interfaces in the C-H model. During each simulation step, the system is evolved into the direction polygon configurations that minimizes the total tissue energy

$E=\underset{\text{Price of (cell) areas}}{\underset{⏟}{\overset{N}{\underset{i=1}{\sum}}\frac{\kappa_{A}}{2}(A_{i}-A_{i}^{0})^{2}}}+\underset{\text{Price of perimeters}}{\underset{⏟}{\overset{N}{\underset{i=1}{\sum}}\frac{\kappa_{P}}{2}(P_{i}-P_{i}^{0})^{2}}}+\underset{\text{Price of junctions}}{\underset{⏟}{2\underset{<\mu,v>}{\sum}\Lambda_{\mu v}l_{\mu v}}}$, (5)

where $\kappa_{A}$ and $\kappa_{P}$ are area and perimeter moduli, respectively. While updating the polygon centroid positions based on the energy minimization, each centroid is further assigned a random motion impetus, which a key aspect in our study of continuous cluster remodeling under impaired PDGFRA^High^ cohesion condition (Fig. 5G-J).

The self-propelled Voronoi model was implemented in Python 3.10. For implementation details and the parameters values used in the simulations in Figs. 5 and S6, as well as additional simulations, see *https://github.com/Gartner-Lab/GutSPV*.

**Random motion of cells**

The break-up of the initial thin layer into distinct clusters requires an impetus to break the initial symmetry: In the continuum model (C-H) we introduce noise into the initial concentration field of the clustering (green) phase, while in the cell-based model (SPV) the motion of the finite-size cells is characterized by random motility sufficient in magnitude to introduce gaps as random events. Note that since in the continuum model the material is transported via diffusion, the model implicitly incorporates the random motion of particles, but at the continuum limit of arbitrary number of such particles the random motion of individual particles is lost in the statistical average of the ensemble. In contrast, in the cell-based model the spatial scale of each individual particle (cell) is sufficiently large such that the random motion of even a single particle can significantly change the geometry. This key difference between the continuum model and its cell-based approximation becomes crucial when investigating whether a given morphogenetic process manifests mainly at a spatial scale of a tissue or a cell: If a process, such as the formation dynamics of large aggregates of cells into clusters, is largely independent of random motion of any individual cell in the aggregate, then the process may be accurately captured by a generalized continuum model that focuses at the dynamics at the scale of whole clusters or tissue, rather than the scale of interaction of individual cells. If, on the other hand, the process is sensitive or dependent on the actions on individual cells, such as the continuous remodeling of clusters in the case of impaired cohesion (Fig. 5H), where the remodeling is characterized by individual cells exchanging clusters in a highly dynamic manner, a cell-level description may be more suitable.

**Conversion between homotypic and heterotypic surface energies**

The SPV model allows for setting surface energies for heterotypic interactions $\sigma_{ij}$ just like in the C-H model, but further also for homotypic interactions $\sigma_{ii}$ that are not present in C-H. While the homotypic parameters $\sigma_{ii}$ allow for concrete expression of the inverse of cell-cell cohesion in the SPV model, we note that in terms of minimizing the total tissue energy they are redundant and can be expressed in terms of adhesive, or heterotypic surface energies. The reduction of the homotypic parameters also allows for direct comparison between SPV and C-H models, since the heterotypic parameters represent analogous surface energies in both models. To reduce the parameters we first compute the new (reduced) heterotypic surface energies for each cell type pair $(i,j)$ as

$\overset{̂}{\sigma_{ij}}=\sigma_{ij}-\frac{\sigma_{ii}+\sigma_{jj}}{2}$,

followed by setting the homotypic energies $\sigma_{ii}$ to zero. Strictly speaking this form of conversion provides only an approximation, as converting the surface energies in this manner affects the total energy of the system in Eq. (5), which may become apparent as a compression (or inflation) of the cells. In practice, however, as long as the absolute magnitudes of the converted energies are relatively modest, the converted system will evolve in close agreement with the non-converted one (see the additional simulations in <https://github.com/Gartner-Lab/GutSPV>).
